# DNMT1 loss leads to hypermethylation of a subset of late replicating domains by DNMT3A

**DOI:** 10.1101/2024.12.19.629414

**Authors:** Ioannis Kafetzopoulos, Francesca Taglini, Hazel Davidson-Smith, Duncan Sproul

## Abstract

Loss of DNA methylation is a hallmark of cancer that is proposed to promote carcinogenesis through gene expression alterations, retrotransposon activation and induction of genomic instability. Cancer-associated hypomethylation does not occur across the whole genome but leads to the formation of partially methylated domains (PMDs). However, the mechanisms underpinning PMD formation remain unclear. PMDs replicate late in S-phase leading to the proposal that they become hypomethylated due to incomplete re-methylation by the maintenance methyltransferase DNMT1 during cell division. Here we investigate the role of DNMT1 in the formation of PMDs in cancer by conducting whole genome bisulfite sequencing (WGBS), repli-seq and ChIP-seq on DNMT1 knockout HCT116 colorectal cancer cells (DNMT1 KO cells). We find that DNMT1 loss leads to preferential hypomethylation in late replicating, heterochromatic PMDs marked by the constitutive heterochromatic mark H3K9me3 or the facultative heterochromatic mark H3K27me3. However, we also observe that a subset of H3K9me3-marked PMDs gain methylation in DNMT1 KO cells. We find that, in DNMT1 KO cells, these hypermethylated PMDs remain late replicating but gain DNMT3A localisation. This is accompanied by loss of heterochromatic H3K9me3 and specific gain of euchromatic H3K36me2. Our observations suggest that hypermethylated PMDs lose their heterochromatic state, enabling their methylation by DNMT3A and the establishment of a hypermethylated, non-PMD state, despite their late replication timing. More generally, our findings suggest that the *de novo* DNMTs play a key role in establishing domain level DNA methylation patterns in cancer cells.

## Introduction

DNA methylation is an epigenetic modification which occurs by the addition of a methyl group on the 5’ position of a cytosine ring. In vertebrates, DNA methylation occurs largely in the context of CpG dinucleotides and the majority of CpGs in the genome are methylated (Lister *et al*, 2009; Suzuki & Bird, 2008). DNA methylation is established in development by the de novo DNA methyltransferases (DNMTs), DNMT3A and DNMT3B (Li *et al*, 1992; Okano *et al*, 1999). Methylation patterns are then maintained predominantly by the maintenance DNA methyltransferase DNMT1 (Dahlet *et al*, 2020), although DNMT3A and B also play a role (Jones & Liang, 2009).

DNA methylation patterns are dysregulated in cancer and overall levels of DNA methylation are reduced in tumours (Feinberg & Vogelstein, 1983; Gama-Sosa *et al*, 1983). This hypomethylation is not uniform across the genome but occurs in interspersed, mega-base scale regions termed partially methylated domains (PMDs) (Berman *et al*, 2011; Hon *et al*, 2012; Lister *et al*, 2011). Cancer-associated hypomethylation is proposed to facilitate tumorigenesis through gene expression alterations, the activation of repetitive elements and the promotion of genome instability (Besselink *et al*, 2023; Howard *et al*, 2008; Wild & Flanagan, 2010).

PMDs feature distinct genomic characteristics compared to the rest of the genome. They have a reduced mean CpG density, are gene poor and the genes within PMDs are generally repressed (Hon *et al*., 2012; Hovestadt *et al*, 2014; Lister *et al*., 2009). This suggests they are heterochromatic in nature. Indeed, PMD chromatin is resistant to nuclease digestion (Spracklin & Pradhan, 2020) and is marked by the repressive histone marks H3K9me3 and H3K27me3 (Hon *et al*., 2012; Salhab *et al*, 2018), associated with constitutive and facultative heterochromatin respectively (Nicetto & Zaret, 2019). In support of the heterochromatic nature of PMDs, they coincide with lamina associated domains (LADs) (Berman *et al*., 2011; Hon *et al*., 2012) and overlap with the HiC defined B-compartment (Salhab *et al*., 2018). In addition, during S-phase, heterochromatin and PMDs are observed to replicate later than euchromatin (Du *et al*, 2019; Salhab *et al*., 2018).

The molecular mechanisms responsible for the formation of PMDs remain unclear. Pulse-chase experiments report that re-methylation of newly synthesised DNA is inefficient and can take several hours following replication (Charlton *et al*, 2018; Ming *et al*, 2020; Stewart-Morgan *et al*, 2023). Re-methylation kinetics are also non-uniform across the genome and heterochromatin is reported to be re-methylated more slowly than euchromatin (Ming *et al*., 2020; Stewart-Morgan *et al*., 2023). This has led to the hypothesis that late replicating regions may not have sufficient time to become re-methylated following DNA replication in rapidly proliferating cancer cells (Petryk *et al*, 2021; Zhou *et al*, 2018). The continuous proliferation of cancer cells could therefore lead to the hypomethylation of heterochromatin and the formation of PMDs with successive cell divisions. Indeed, PMD methylation levels negatively correlate with the number of mutations in tumours (Zhou *et al*., 2018) suggesting that PMDs lose DNA methylation as a function of the number of cell divisions that have occurred. Given that DNMT1 is primarily responsible for maintaining DNA methylation following replication (Petryk *et al*., 2021), this model predicts a central role for DNMT1 in the formation of PMDs. In support of this hypothesis, DNMT1 has been observed to be excluded from heterochromatic, late replicating satellite DNA in senescent cells where extensive hypomethylation of PMDs occurs (Cruickshank *et al*, 2013).

To understand how maintenance methylation in cancer cells varies with replication timing and chromatin structure, we have interrogated DNA methylation changes in DNMT1 knock-out HCT116 cells (DNMT1 KO). While we observe that late-replicating, heterochromatic regions lose more DNA methylation in DNMT1 KO cells than early replicating, euchromatic regions, we also observe that a subset of H3K9me3-marked late-replicating regions gain DNA methylation. This associates with the recruitment of DNMT3A, loss of H3K9me3 and specific gain of H3K36me2. These results therefore suggest that DNMT3A plays an important role in counteracting hypomethylation in PMDs.

## Results

### Ablation of DNMT1 leads to preferential hypomethylation of partially methylated domains

To understand the role of DNMT1 in shaping the cancer epigenome at the domain level, we compared the methylomes of HCT116 cells to their DNMT1 knock-out (DNMT1 KO cells) derivatives (Rhee *et al*, 2000) using Whole Genome Bisulfite Sequencing (WGBS) (HCT116 mean coverage= 1.6x, DNMT1 KO mean coverage= 2.4x). As complete removal of DNMT1 in somatic cells is lethal, DNMT1 KO cells express low levels of a truncated DNMT1 (Egger *et al*, 2006; Spada *et al*, 2007).

Before examining the nature of the changes in DNA methylation in DNMT1 KO cells, we first assessed the DNA methylation landscape in HCT116 cells. We used the WGBS to quantify mean DNA methylation levels in 10Kb bins across the genome, finding that they were bimodally distributed due to the presence of PMDs (*Figure 1A-B*). To understand the properties of these PMDs, we used methpipe to define 546 HCT116 PMDs on autosomal chromosomes (Decato *et al*, 2020). We then defined the rest of the genome as 556 HCT116 highly methylated domains (HMDs, see methods for details).

**Figure 1.**
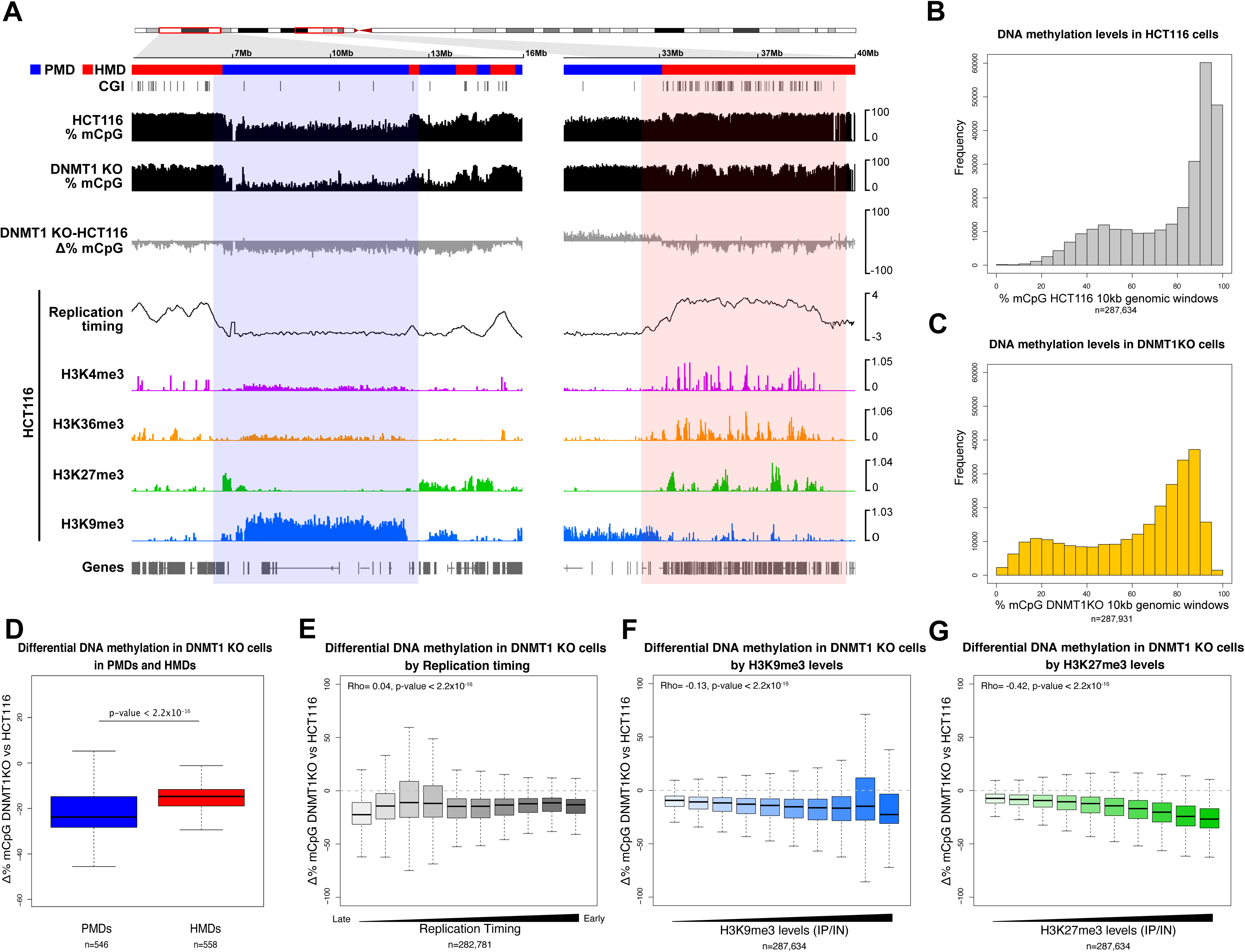
Ablation of DNMT1 leads to preferential hypomethylation of partially methylated domains. (**A**) Representative genomic loci showing changes of DNA methylation at a PMD and HMD in DNMT1 KO cells. Genome browser plots showing absolute (black) and differential (grey) DNA methylation levels (mCpG) alongside HCT116 histone modification ChIP-seq and repli-seq. DNA methylation levels are plotted in 10 kb genomic windows. ChIP-seq tracks are normalised log_10_ IP/IN. Replication timing data are loess smoothed repli-seq early/late ratios over 10 kb. Representative PMD and HMD are indicated by the coloured boxes. CGI = CpG island. (**B, C**) Histograms of DNA methylation levels in HCT116 (**B**) and DNMT1 KO (**C**) cells. Mean DNA methylation levels were calculated for 10 kb genomic windows. (**D**) Boxplot showing differential DNA methylation between DNMT1 KO and HCT116 at PMDs (n = 546 domains) and HMDs (n = 558 domains) defined in HCT116 cells. P-value from two-sided Wilcoxon rank sum test. (**E**) Boxplot showing difference in DNA methylation between DNMT1 KO and HCT116 cells in 10 kb genomic windows divided in deciles according to their HCT116 replication timing. Replication timing data are loess smoothed repli-seq early/late ratios over 10 kb. (**F**) Boxplot showing difference in DNA methylation between DNMT1 KO and HCT116 cells in 10 kb genomic windows divided in deciles according to their H3K9me3 levels in HCT116 cells. ChIP-seq data are normalised IP/IN. (**G**) Boxplot showing difference in DNA methylation between DNMT1 KO and HCT116 cells in 10 kb genomic windows divided in deciles according to their H3K27me3 levels in HCT116 cells. ChIP-seq data are normalised IP/IN. For boxplots: Lines = median; box = 25th–75th percentile; whiskers = 1.5 × interquartile range from box. All histone ChIP-seq and repli-seq data shown are derived from the mean of two biological replicates. In (**E-G**), n is the number of windows analysed shown below the plots and the Spearman’s correlation coefficient (Rho) is shown alongside its associated p-value.

PMDs have previously been reported to replicate late in S-phase and be marked with heterochromatic histone modifications (Berman *et al*., 2011; Hon *et al*., 2012). To understand whether this was also the case in HCT116 cells, we performed repli-seq to measure replication timing alongside ChIP-seq for the histone modifications H3K4me3, H3K9me3, H3K27me3 and H3K36me3. We then compared replication timing between HCT116 PMDs and HMDs confirming that the replication timing of PMDs was significantly later than that of HMDs (*Figure S1A*, p < 2.2×10^-16^, Wilcoxon rank sum test). Both the constitutive heterochromatin-associated H3K9me3 and the facultative heterochromatin-associated H3K27me3 marks had significantly higher levels in HCT116 PMDs compared to HMDs (*Figure S1B-C*, both p < 2.2×10^-16^, Wilcoxon rank sum tests). H3K4me3 and H3K36me3 were observed in peaks around promoters and gene bodies in HMDs respectively (*Figure S1D*). However, we also noted a low, broad enrichment of both marks in PMDs (*Figure 1A*).

Further investigation revealed that HCT116 PMDs were split into 253 predominantly marked by H3K9me3 and 293 predominantly marked by H3K27me3 (*Figure S1E*). H3K9me3-marked PMDs were bordered by H3K27me3 (*Figure 1A, S1E*), a pattern previously reported in IMR90 and HCC1954 cells (Hon *et al*., 2012). The median length of H3K9me3-enriched PMDs was 1.86Mb compared to 0.59Mb for H3K27me3-enriched PMDs. In addition, the mean methylation level of H3K9me3-enriched PMDs was significantly lower than that of H3K27me3 enriched PMDs (*Figure S1F*, p = 6.85×10^-13^, Wilcoxon rank sum test) and the mean replication timing of H3K9me3-enriched PMDs was significantly later than H3K27me3-enriched PMDs (*Figure S1G*, p < 2.2×10^-16^, Wilcoxon rank sum test).

We then asked how the methylome changed in DNMT1 KO cells. Global mean methylation levels measured by WGBS in DNMT1 KO cells were 59.2% as compared to 75.7% in HCT116 cells (*Figure S1H*), a similar reduction to previous reports (Rhee *et al*., 2000). Despite this global reduction, mean methylation levels across the genome in DNMT1 KO cells had a bimodal distribution like that observed in HCT116 cells (*Figure 1B-C*).

To understand whether different genomic domains might display differential requirement for DNMT1, we then compared the degree of methylation loss observed in DNMT1 KO cells between HCT116 PMDs and HMDs. While both PMDs and HMDs had reduced methylation in DNMT1 KO cells, the level of methylation loss in PMDs observed in DNMT1 KO cells was significantly greater than in HMDs (*Figure 1D*, p < 2.2×10^-16^, Wilcoxon rank sum test).

To understand how these differences in the degree of methylation loss related to the properties of PMDs, we then analysed how mean methylation changes across the genome related to replication timing and PMD-associated histone marks in HCT116 cells. Despite the general late replication of PMDs, we observed a weak correlation between DNA methylation reduction and HCT116 replication timing measured by repli-seq (*Figure 1E*, Rho = 0.048, p < 2.2×10^-16^, Spearman’s rank correlation). In contrast we observed stronger, significant correlations between mean H3K9me3 or H3K27me3 levels and DNA methylation reduction in DNMT1 KO cells (*Figures 1F-G*, Rho = −0.136 and Rho = −0.428 respectively, both p < 2.2×10^-16^, Spearman’s rank correlations).

Taken together, these analyses suggest that the loss of methylation observed upon removal of DNMT1 in cancer cells is non-uniform and DNA methylation is preferentially lost from PMDs associated with the histone marks H3K27me3 and H3K9me3.

### A subset of H3K9me3-marked PMDs are hypermethylated in DNMT1 knockout cells

When examining differences in the methylation levels of PMDs and HMDs between HCT116 and DNMT1 KO cells, we noted that a subset of PMDs had higher levels of DNA methylation in DNMT1 KO cells (*Figure 2A-B*). This was surprising given the reduced function of the major maintenance DNMT in these cells and the bias towards greater methylation losses in PMDs in DNMT1 KO cells.

**Figure 2.**
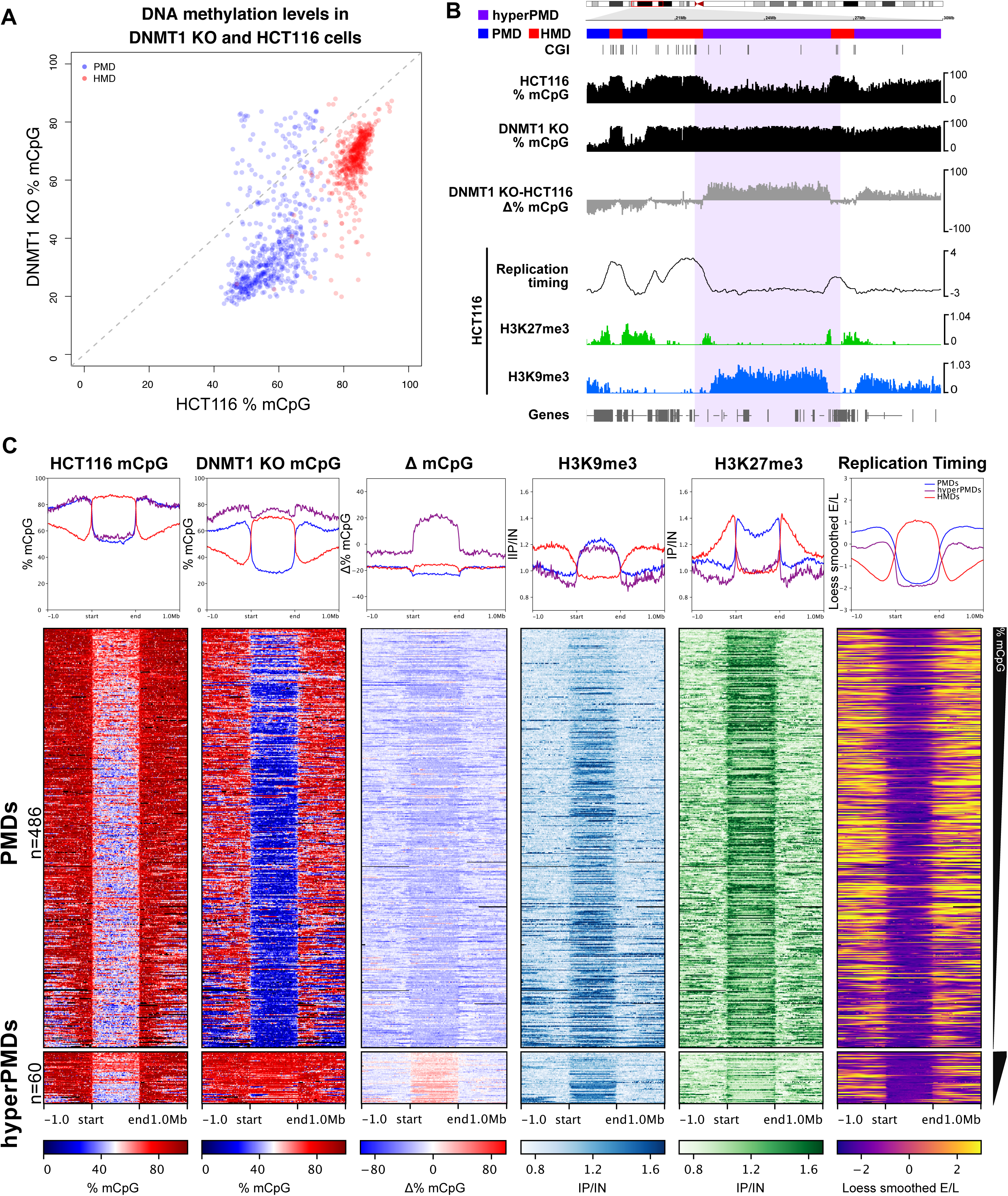
A subset of H3K9me3-marked PMDs are hypermethylated in DNMT1 knockout cells. (**A**) Scatter plot of mean methylation levels at PMDs and HMDs in DNMT1 KO cells versus HCT116 cells. (**B**) Representative genomic loci showing changes of DNA methylation at a hypermethylated PMD in DNMT1 KO cells. Genome browser plots showing absolute (black) and differential (grey) DNA methylation levels (mCpG) alongside HCT116 histone modification ChIP-seq and repli-seq. DNA methylation levels are plotted in 10 kb genomic windows. ChIP-seq tracks are normalised log_10_ IP/IN. Replication timing data are loess smoothed repli-seq early/late ratios over 10 kb. Representative hypermethylated PMD and HMD are indicated by the coloured boxes. CGI = CpG islands. (**C**) Heatmaps and pileup plots of HCT116 H3K9me3, H3K27me3 and DNA methylation levels alongside replication timing for hypermethylated PMDs (n= 60) and all other HCT116 PMDs (n= 486). ChIP-seq data are mean normalised IP/IN. DNA methylation levels are mean % mCpG. Replication timing data are mean loess smoothed repli-seq early/late ratios over 10kb. PMDs are aligned and scaled to the start and end points of each domain and ranked based on their mean methylation levels in HCT116 cells. All histone ChIP-seq and repli-seq data shown are derived from the mean of two biological replicates.

To understand the factors that led to this increase in methylation at some PMDs upon removal of DNMT1, we defined hypermethylated PMDs as those where mean methylation levels in DNMT1 KO cells were ≥ 5% higher than in HCT116 cells (60 out of 546 PMDs, 10.9%) (*Figure 2B, S2A*). Gains of methylation were significantly more likely to occur at PMDs than HMDs and only 3 out of 556 HMDs had ≥ 5% mean methylation in DNMT1 KO cells compared to HCT116 cells (p = 6.33×10^-16^, Fisher’s exact test). These hypermethylated PMDs were distributed across all the autosomal chromosomes and were longer than all other HCT116 PMDs (*Figure S2B*, median length 2.12Mb versus 0.94Mb of all other PMDs, p = 1.52×10^-5^, Wilcoxon rank sum test).

We then asked whether hypermethylated PMDs were associated with a distinct set of chromatin marks compared to all other HCT116 PMDs. Hypermethylated PMDs had similar levels of H3K9me3 but lower levels of H3K27me3 than all other HCT116 PMDs (*Figures 2B-C, S2C-D*, p = 0.59 and < 2.2×10^-16^ respectively, Wilcoxon rank sum tests). Hypermethylated PMDs also replicated slightly but significantly later than all other HCT116 PMDs (*Figure 2B-C, S2E*). Consistent with these characteristics, we found that the hypermethylated PMDs were significantly enriched in H3K9me3-marked PMDs defined in our previous analysis (43 out of 60 hypermethylated PMDs were H3K9me3-marked PMDs, 71.66%, p = 2.42×10^-5^, Fisher’s exact test).

These observations suggest a subset of H3K9me3-marked PMDs have higher levels of methylation in DNMT1 KO cells despite their severe reduction of maintenance methyltransferase activity.

### The replication timing of hypermethylated PMDs remains similar in DNMT1 KO cells

We next sought to understand the factors that could have led to the higher levels of methylation observed at hypermethylated PMDs in DNMT1 KO cells. The late replication of PMDs in S-phase is proposed to underpin their loss of methylation with successive cell divisions (Aran *et al*, 2011; Salhab *et al*., 2018; Zhou *et al*., 2018). We therefore wondered whether the higher methylation levels of hypermethylated PMDs was linked to a shift in their replication timing.

We compared the replication programs of DNMT1 KO and HCT116 cells using repli-seq (*Figure 3A*). The overall replication timing of DNMT1 KO cells was highly significantly correlated to that of HCT116 cells (*Figure S3A*, Rho = 0.91, p < 2.2×10^-16^, Spearman’s rank correlation). As in HCT116 cells, DNMT1 KO cell replication timing was significantly but weakly correlated with methylation levels consistent with highly methylated parts of the DNMT1 KO genome having earlier replication timing than lowly methylated regions (*Figure S3B-C*, HCT116: Rho = 0.47, DNMT1 KO: Rho = 0.27, both p < 2.2×10^-16^, Spearman’s rank correlations). Similar to our observations in HCT116 cells, HCT116 PMDs were significantly later replicating than HCT116 HMDs in DNMT1 KO (*Figure S1A, S3D*, p < 2.2×10^-16^, Wilcoxon rank sum test).

**Figure 3.**
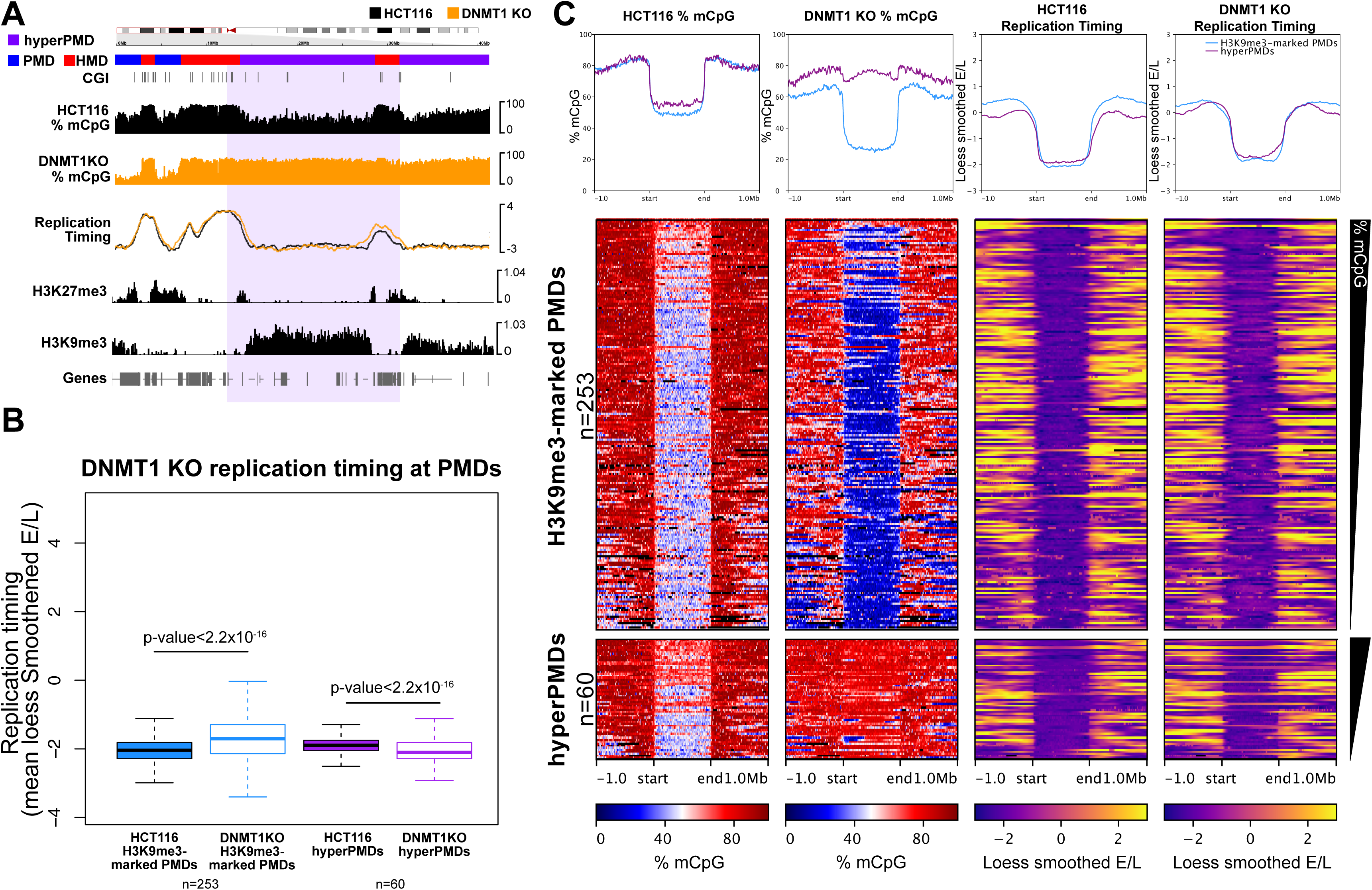
The replication timing of hypermethylated PMDs remains similar in DNMT1 KO cells. (**A**) Representative genomic loci showing DNA methylation and replication timing HCT116 and DNMT1 KO cells. Genome browser plots showing DNA methylation levels (mCpG) alongside HCT116 histone modification ChIP-seq and repli-seq. DNA methylation levels are plotted in 10 kb genomic windows. ChIP-seq tracks are normalised log_10_ IP/IN. Replication timing data are loess smoothed repli-seq early/late ratios over 10 kb. Representative hypermethylated PMD is indicated by the coloured box. CGI = CpG islands. (**B**) Boxplot comparing HCT116 and DNMT1 KO replication timing at hypermethylated PMDs (n = 60) and other H3K9me3-marked PMDs (n = 253 Domains). Replication timing data are mean loess smoothed repli-seq early/late ratios over 10kb. Box plot annotation: lines = median; box = 25th–75th percentile; whiskers = 1.5 × interquartile range from box. P-values are from two-sided Wilcoxon rank sum tests. (**C**) Heatmaps and pileup plots of HCT116 and DNMT1 KO DNA methylation levels alongside replication timing for hypermethylated PMDs (n= 60) and all other H3K9me3-marked PMDs (n= 253). ChIP-seq data are mean normalised IP/IN. DNA methylation levels are mean % mCpG. Replication timing data are mean loess smoothed repli-seq early/late ratios over 10kb. PMDs are aligned and scaled to the start and end points of each domain and ranked based on their mean methylation levels in HCT116 cells. All histone ChIP-seq and repli-seq data shown are derived from the mean of two biological replicates.

We then focused specifically on understanding whether the replication timing of hypermethylated PMDs changed in DNMT1 KO cells compared to other HCT116 H3K9me3-marked PMDs. Despite their high levels of methylation in DNMT1 KO cells, overall, hypermethylated PMDs remained late replicating in DNMT1 KO cells (*Figure 3A-C*). In fact, hypermethylated PMDs were observed to replicate slightly but significantly later in DNMT1 KO cells compared to HCT116 cells (*Figure 3B*, p < 2.2×10^-16^, Wilcoxon rank sum test). In contrast, other H3K9me3-enriched HCT116 PMDs replicated slightly but significantly earlier in DNMT1 KO cells than HCT116 cells (*Figure 3B*, p < 2.2×10^-16^, Wilcoxon rank sum test).

Overall, these analyses suggest that despite the global associations between late replication timing and low levels of methylation, hypermethylated PMDs do not become earlier replicating in DNMT1 KO cells.

### DNMT3A localises to hypermethylated PMDs

We next asked whether the higher methylation levels at hyper PMDs in DNMT1 KO cells might result from activity of the *de novo* methyltransferases, DNMT3A or 3B.

To understand whether this might be the case, we first examined whether DNMT3A or 3B were differentially expressed between HCT116 and DNMT1 KO cells using RNA-seq data. Consistent with the knockout of DNMT1, DNMT1 RNA levels were significantly reduced in DNMT1 KO cells compared to HCT116 cells (*Figure S4A*, 2.88-fold, p = 1.59×10^-10^). However, only small, non-significant differences in the RNA levels of both DNMT3A and 3B were observed in DNMT1 KO cells relative to HCT116 cells (*Figure S4A*). This suggests DNMT3A and B levels were not greatly altered in DNMT1 KO cells.

We then asked whether DNMT3A or 3B localised differently in HCT116 and DNMT1 KO cells by expressing T7-tagged DNMT3A or 3B in HCT116 and DNMT1 KO cells and conducting ChIP-seq to determine their localisation. For this experiment we selected the main catalytically active versions of the enzymes found in HCT116 cells, DNMT3A1 and DNMT3B2 (Masalmeh *et al*, 2021; Taglini *et al*, 2024). We have also previously shown that ectopically expressed DNMT3B recapitulates the localisation of endogenous DNMT3B in HCT116 cells (Taglini *et al*., 2024).

In HCT116 cells, both DNMT3A and 3B levels were significantly higher in HMDs compared to PMDs (*Figure S4B-D*). HCT116 DNMT3B ChIP-seq signal was significantly correlated with H3K36me3 ChIP-seq signal (*Figure S4E*, Rho = 0.54, p < 2.2×10^-16^, Spearman’s rank correlation) whereas HCT116 DNMT3A ChIP-seq signal showed lower correlations with H3K36me3 (*Figure S4F*, Rho = 0.12, p < 2.2×10^-16^, Spearman’s rank correlation), as previously reported (Baubec et al., 2015; Weinberg et al., 2019).

We next asked how DNMT3A and 3B localised in DNMT1 KO cells as compared to HCT116 cells. The overall profiles of DNMT3A and 3B in DNMT1 KO cells were equivalent to that observed in HCT116 cells with significantly more of both enzymes observed in HMDs compared to PMDs (*Figure S4B-D*). In addition, both DNMT3A and 3B ChIP-seq signals across the genome were significantly correlated between HCT116 and DNMT1 KO cells (*Figure S4G-H*, both Rho = 0.64, p < 2.2×10^-16^, Spearman’s rank correlations).

To understand how DNMT3A and 3B might contribute to the higher levels of methylation at hypermethylated PMDs in DNMT1 KO cells, we examined their localisation in these domains. In HCT116 cells, levels of both enzymes in hypermethylated PMDs were low and equivalent to other HCT116 H3K9me3-marked PMDs (*Figure 4A-D*). Both enzymes were depleted from HCT116 H3K9me3-marked PMDs in DNMT1 KO cells, but DNMT3A levels were significantly increased at hypermethylated PMDs in DNMT1 KO cells when compared to HCT116 cells (*Figure 4A-B, D*, p < 2.2×10^-16^, Wilcoxon rank sum test). Despite this increase, DNMT3A levels at hypermethylated PMDs in DNMT1 KO cells were lower than was observed in the surrounding HMD regions in DNMT1 KO cells (*Figure 4A, S4B, D*). DNMT3B levels were not significantly increased at hypermethylated PMDs in DNMT1 KO cells (*Figure 4A, C-D*, p=0.83, Wilcoxon rank sum test) but DNMT3B levels at other H3K9me3-marked PMDs were significantly reduced in DNMT1 KO cells compared to HCT116 cells (*Figure 4C, p <* 2.2×10^-16^). This suggests that the higher levels of DNA methylation at hypermethylated PMDs in DNMT1 KO cells are associated with DNMT3A localisation.

**Figure 4.**
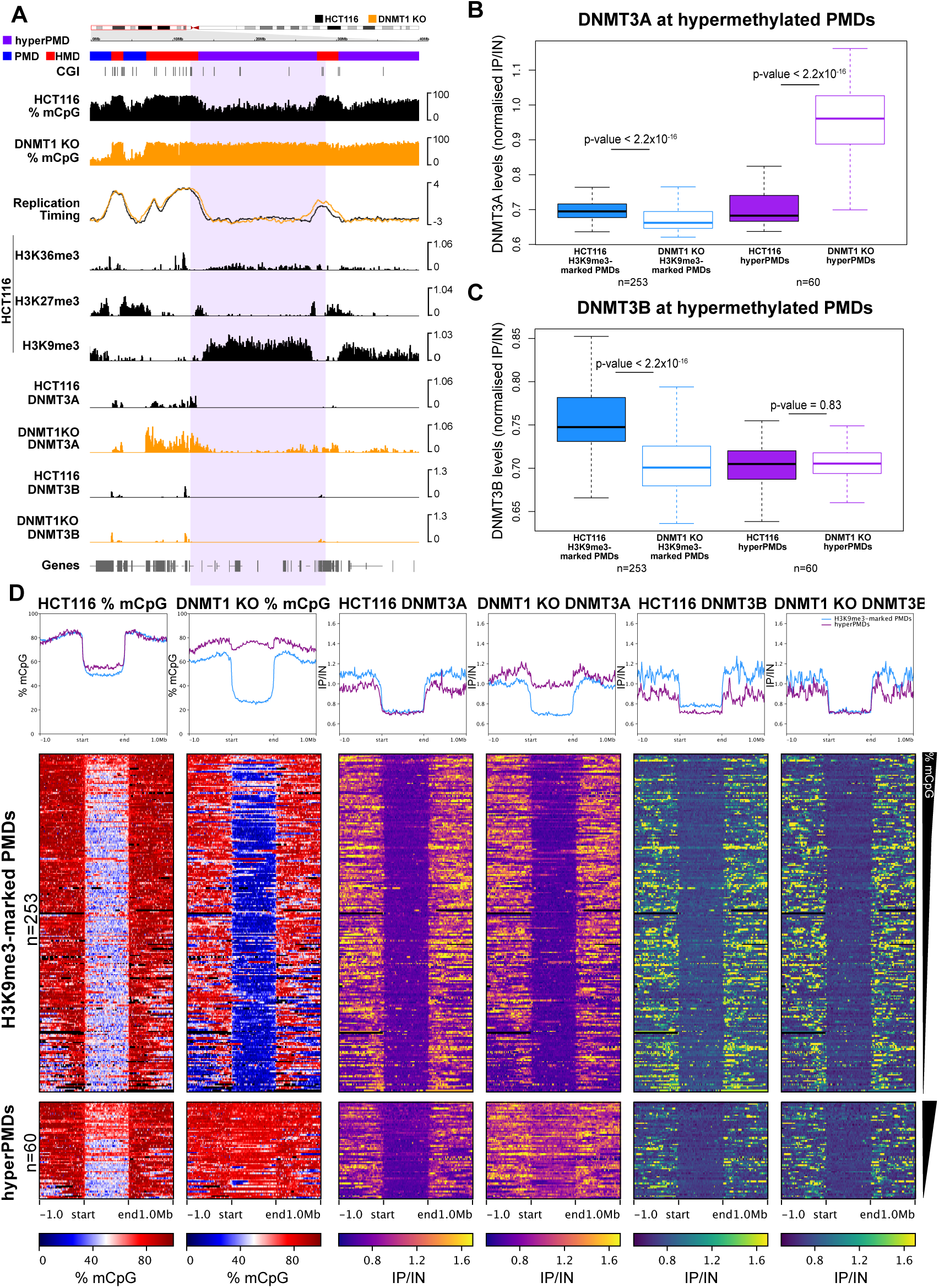
DNMT3A localises to hypermethylated PMDs. (**A**) Representative genomic loci showing DNMT3A localisation at a hypermethylated PMD in DNMT1 KO cells. Genome browser plots showing DNA methylation levels (mCpG) alongside DNMT3A/B ChIP-seq and HCT116 histone modifications and repli-seq. DNA methylation levels are plotted in 10 kb genomic windows. ChIP-seq tracks are normalised log_10_ IP/IN. Replication timing data are loess smoothed repli-seq early/late ratios over 10 kb. Representative hypermethylated PMD is indicated by the coloured box. CGI = CpG islands. (**B**) Boxplot showing DNMT3A levels at hypermethylated PMDs (n = 60 domains) compared to other H3K9me3-marked PMDs (n = 253 domains). ChIP-seq data are mean normalised IP/IN. (**C**) Boxplot showing DNMT3B levels at hypermethylated PMDs (n = 60 domains) compared to other H3K9me3-marked PMDs (n = 253 Domains). ChIP-seq data are mean normalised IP/IN. (**D**) Heatmaps and pileup plots of HCT116 and DNMT1 KO DNA methylation levels alongside DNMT3A/B for hypermethylated PMDs (n= 60) and all other H3K9me3-marked PMDs (n= 253). ChIP-seq data are mean normalised IP/IN. DNA methylation levels are mean % mCpG. PMDs are aligned and scaled to the start and end points of each domain and ranked based on their mean methylation levels in HCT116 cells. For boxplots: Lines = median; box = 25th–75th percentile; whiskers = 1.5 × interquartile range from box. All p-values are from two-sided Wilcoxon rank sum tests. All histone ChIP-seq and repli-seq data shown are derived from the mean of two biological replicates.

### Hypermethylated PMDs lose H3K9me3 and gain H3K36me2

We next sought to understand why DNMT3A was recruited to hypermethylated PMDs in DNMT1 KO cells. The recruitment of DNMTs to the genome is largely driven by the recognition of histone marks (Tibben & Rothbart, 2024). However, DNMT3A has also previously been reported to be excluded from heterochromatin (Stroud *et al*, 2017; Yamanaka *et al*, 2019).

We therefore conducted ChIP-seq for the heterochromatic histone marks H3K9me3 and H3K27me3 in DNMT1 KO cells and compared their profile to HCT116 cells. The overall profiles of H3K9me3 and H3K27me3 were significantly correlated between the two cell types (*Figure S5A-C*, Rho = 0.56 and 0.71 respectively, both p < 2.2×10^-16^, Spearman’s rank correlations). In support of a broadly similar genome-wide localisation of the two marks, Hidden Markov Model defined H3K9me3 and H3K27me3 domains in the two cell types also significantly overlapped (H3K9me3: Jaccard = 0.44, Fisher’s test p < 2.2×10^-16^, H3K27me3 Jaccard = 0.57, Fisher’s test p < 2.2×10^-16^). However, we noted that levels of H3K9me3 in H3K9me3-marked PMDs were slightly reduced and occasional spreading of the H3K9me3 mark was observed (*Figure 5A, S5A*).

**Figure 5.**
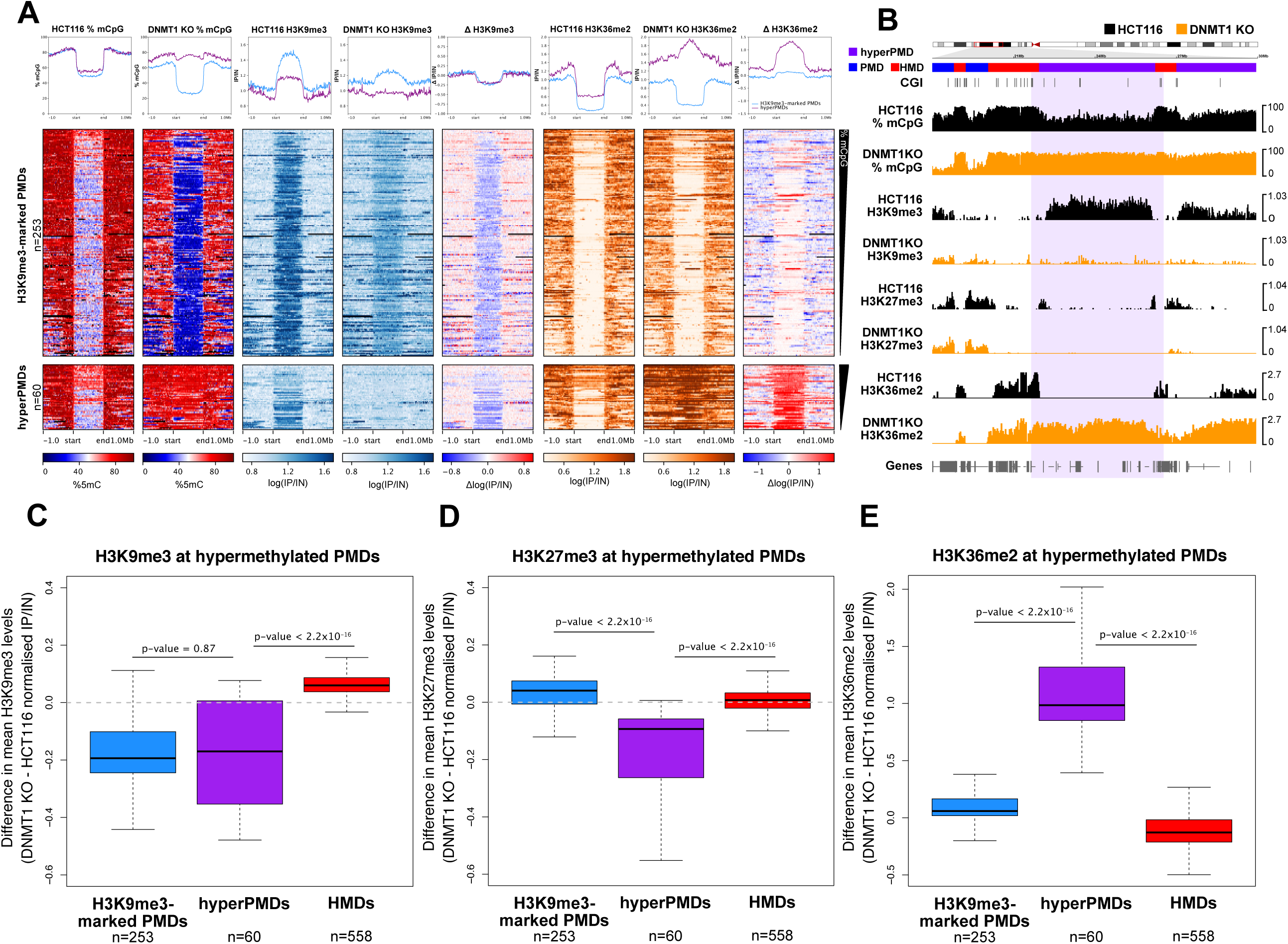
Hypermethylated PMDs lose H3K9me3 and gain H3K36me2. (**A**) Heatmaps and pileup plots of HCT116 and DNMT1 KO DNA methylation levels alongside H3K9me3 and H3K36me2 levels for hypermethylated PMDs (n= 60) and all other H3K9me3-marked PMDs (n= 253). ChIP-seq data are mean normalised IP/IN. DNA methylation levels are mean % mCpG. PMDs are aligned and scaled to the start and end points of each domain and ranked based on their mean methylation levels in HCT116 cells. (**B**) Representative genomic loci showing H3K9me3 and H3K36me2 levels at a hypermethylated PMD in DNMT1 KO cells. Genome browser plots showing DNA methylation levels (mCpG) alongside H3K9me3 and H3K36me2 ChIP-seq. DNA methylation levels are plotted in 10 kb genomic windows. ChIP-seq tracks are normalised log_10_ IP/IN. Representative hypermethylated PMD is indicated by the coloured box. CGI = CpG islands. (**C**) Boxplot showing change in H3K9me3 levels at H3K9me3-marked PMDs (n = 253 domains), hypermethylated PMDs (n = 60 domains) and HMDs (n = 558 domains). ChIP-seq data are difference in mean normalised IP/IN between DNMT1 KO and HCT116 cells. (**D**) Boxplot showing change in H3K27me3 levels at H3K9me3-marked PMDs (n = 253 domains), hypermethylated PMDs (n = 60 domains) and HMDs (n = 558 domains). ChIP-seq data are difference in mean normalised IP/IN between DNMT1 KO and HCT116 cells. (**E**) Boxplot showing change in H3K36me2 levels at H3K9me3-marked PMDs (n = 253 domains), hypermethylated PMDs (n = 60 domains) and HMDs (n = 558 domains). ChIP-seq data are difference in mean normalised IP/IN between DNMT1 KO and HCT116 cells. For boxplots: Lines = median; box = 25th–75th percentile; whiskers = 1.5 × interquartile range from box. All p-values are from two-sided Wilcoxon rank sum tests. All histone ChIP-seq data shown are derived from the mean of two biological replicates.

We then went on to specifically ask whether H3K9me3 or H3K27me3 were altered at hypermethylated PMDs in DNMT1 KO cells. We observed that hypermethylated PMDs showed a significant decrease in both H3K9me3 and H3K27me3 levels in DNMT1 KO cells compared to HCT116 cells (*Figure 5A-D and S5D*, p = 2.06×10^-6^, Wilcoxon rank sum test). We also observed a decrease in H3K9me3 levels from H3K9me3-marked PMDs in DNMT1 KO cells (*Figures 5C and S5B)*. However, levels of H3K9me3 in hypermethylated PMDs decreased to that of background while levels at other H3K9me3-marked PMDs remained above background. This was supported by the observation that hypermethylated PMDs were significantly less likely to overlap H3K9me3 domains in DNMT1 KO cells than other H3K9me3-marked PMDs (11 of 60, 18.33%, hypermethylated PMDs overlap DNMT1 KO H3K9me3 domains whereas 192 of 210, 91.42%, of HCT116 H3K9me3-marked PMDs do, Fisher’s test p < 2.2×10^-16^). Hypermethylated PMDs also showed a significant decrease in H3K27me3 levels in DNMT1 KO cells as compared to HCT116 cells (*Figure 5A, D, S5D,* p = 1.71×10^-10^, Wilcoxon rank sum test). This reflected the loss of H3K27me3 marking the borders of hypermethylated PMDs in HCT116 (*Figure S5D*). Overall, this suggests that hypermethylated PMDs lose heterochromatic histone marks in DNMT1 KO cells.

DNMT3A also localises in the genome through the association of its PWWP domain with H3K36me2 (Weinberg et al., 2019). To understand whether differences in H3K36me2 might be involved in the localisation of DNMT3A to hypermethylated PMDs, we therefore performed ChIP-seq for H3K36me2 in HCT116 and DNMT1 KO cells. The distribution of H3K36me2 in DNMT1 KO cells was significantly correlated with that in HCT116 cells (*Figure S5E*, Rho= 0.68, p < 2.2×10^-16^, Spearman’s rank correlation) and in both HCT116 and DNMT1 KO cells we observed that H3K36me2 was primarily associated with euchromatic HMD regions and depleted from PMDs (*Figure S5A*).

We then specifically examined H3K36me2 levels at hypermethylated PMDs. Like other H3K9me3-marked and H3K27me3-marked PMDs, hypermethylated-PMDs had low levels of H3K36me2 in HCT116 cells (*Figure 5A-B*). In DNMT1 KO cells, H3K36me2 levels remained low in both H3K9me3-marked and H3K27me3-marked PMDs (*Figure 5A-5B, S5H*). However, hypermethylated PMDs specifically showed significant increases of H3K36me2 in DNMT1 KO cells compared to HCT116 cells (*Figure 5A-B, E,* p < 2.2×10^-16^, Wilcoxon sum test).

Taken together, this suggests that a subset of H3K9me3-marked PMDs from HCT116 cells lose this mark in DNMT1 KO cells in association with gain of H3K36me2, leading to the recruitment of DNMT3A and subsequent higher levels of DNA methylation.

## Discussion

Here we have analysed the effect of DNMT1 knockout on PMDs in colorectal cancer cells. We find that losses of methylation in DNMT1 KO cells are biased towards PMDs. However, we also observe that several H3K9me3-marked regions have increased levels of methylation in these cells and lose PMD identity. In association with this, these hypermethylated PMDs lose H3K9me3, gain H3K36me2 and recruit the *de novo* methyltransferase DNMT3A (*Figure 6*). These observations suggest a key role for *de novo* methyltransferase activity in determining which parts of the genome become PMDs in cancer. We propose that the activity of DNMT1 is not sufficient to maintain DNA methylation levels throughout the genome and that re-iterative activity of DNMT3A and B prevents euchromatic regions becoming hypomethylated. In contrast, a lack of this *de novo* activity leads to the formation of PMDs in heterochromatic regions in tumours.

**Figure 6.**
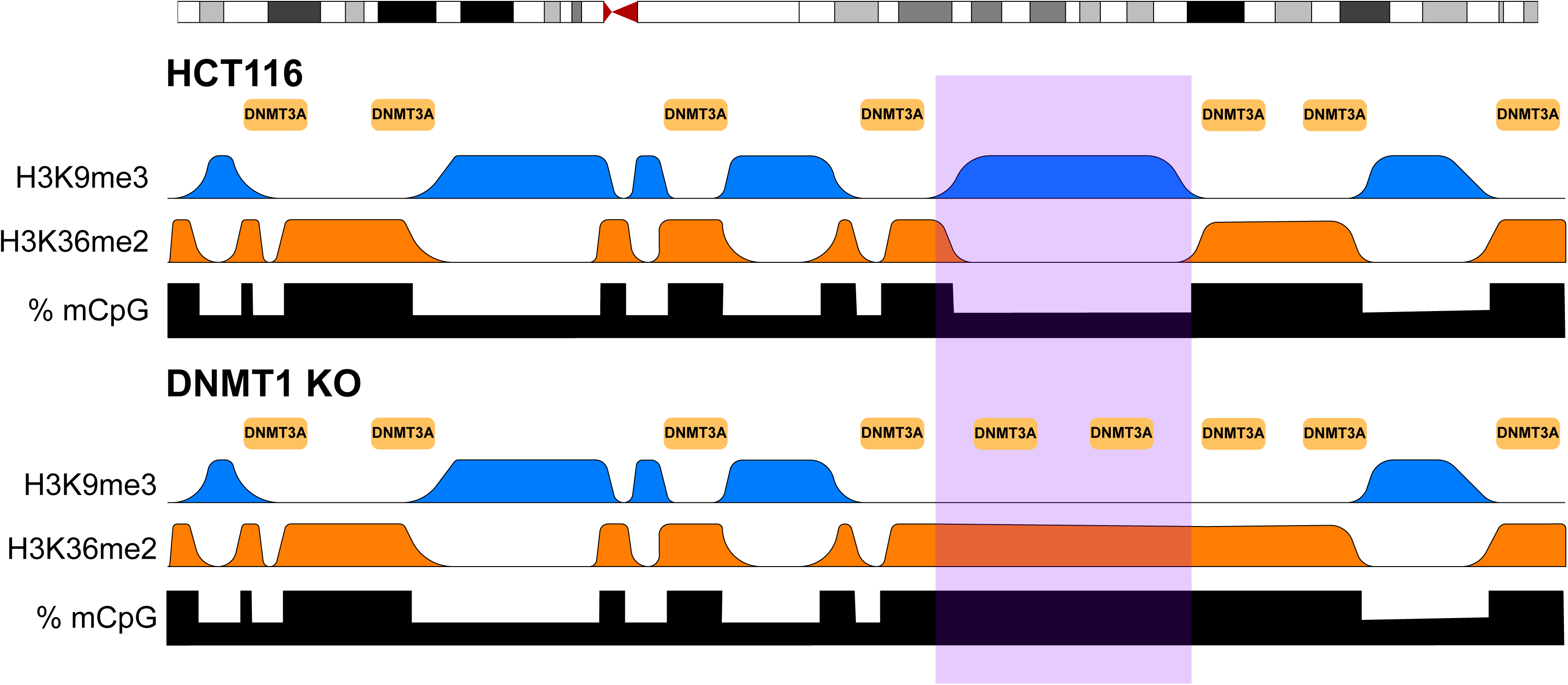
DNMT1 loss leads to hypermethylation of a subset of late replicating domains by DNMT3A. Schematic showing distribution of DNA methylation, histone modifications and DNMT3A at the domain level in HCT116 and DNMT1 KO cells. DNMT3A predominantly localises to H3K36me2-marked regions through the action of its PWWP domain (Weinberg *et al*., 2019) and is excluded from heterochromatic domains. We propose that chromatin remodelling in DNMT1 KO cells leads to loss of heterochromatic H3K9me3 in some regions and gain of H3K36me2. This causes localisation of DNMT3A and gains of DNA methylation.

Previous work has connected the hypomethylation of PMDs to their late replication in S-phase (Zhou *et al*., 2018). Analysis of newly replicated DNA by bisulfite sequencing suggests that re-methylation can take several hours (Charlton *et al*., 2018) and may occur in two phases (Ming *et al*., 2020). This is supported by mass-spectrometry analysis suggesting that some re-methylation occurs post-mitosis (Stewart-Morgan *et al*., 2023). These studies have led to the hypothesis that there is insufficient time to maintain DNA methylation patterns in late replicating regions in rapidly dividing cancer cells resulting in their hypomethylation. Here we do observe that DNMT1 knockout in a colorectal cancer cell line biases DNA methylation losses to PMDs which are late replicating. This parallels a recent study reporting that losses of DNA methylation upon shRNA depletion of DNMT1 in HCT116 cells are biased to low-CpG density regions that are enriched in PMDs (Tiedemann *et al*, 2024). However, we observe weak correlations between DNA methylation loss and replication timing in DNMT1 KO cells. Furthermore, we observe that some regions of the genome gain DNA methylation in these cells while remaining late replicating. Overall, our results suggest that the formation of PMDs is not tightly linked to the replication timing program.

We instead observe stronger correlations between methylation levels and localisation of the *de novo* DNMTs. Normally DNMT3A localises to H3K36me2, a mark broadly distributed in euchromatin (Weinberg *et al*, 2019) whereas DNMT3B is localised to the transcription-associated mark H3K36me3 in mouse embryonic stem cells (Baubec *et al*, 2015) and colorectal cancer cells (Masalmeh *et al*., 2021). It is possible that the activity of these *de novo* methyltransferases compensates for any failure to maintain methylation at euchromatin by DNMT1. Normally such activity is low at heterochromatin and this inability to correct errors made by DNMT1 could underpin the formation of PMDs. This hypothesis is supported by our observation that DNMT3A localises to hypermethylated PMDs in DNMT1 KO cells and a report that *Kras* mutant mouse lung cancers lacking DNMT3A have a more uniformly hypomethylated genome than those possessing DNMT3A (Raddatz *et al*, 2012). Previously, we also observed that PWWP domain mutations can cause DNMT3B to re-localise to H3K9me3-marked heterochromatin also leading to the hypermethylation of PMDs (Taglini *et al*., 2024). Our observations suggest that previously reported differences in the re-methylation kinetics of different parts of the genome following DNA replication (Charlton *et al*., 2018; Ming *et al*., 2020; Stewart-Morgan *et al*., 2023), could be caused by varying levels of *de novo* activity rather than variation in the kinetics by re-methylation of DNMT1 across the genome.

We observe that the localisation of DNMT3A to hypermethylated PMDs is associated with both a loss of H3K9me3 and a gain of H3K36me2. Previous work has observed that H3K9me3-marked heterochromatin excludes DNMT3A in brain development (Stroud *et al*., 2017). In sperm development, heterochromatic regions gain DNA methylation upon loss of H3K9me3 and gain of H3K36me2 (Shirane *et al*, 2020; Yamanaka *et al*., 2019) suggesting that both marks contribute to DNMT3A localisation (Janssen & Lorincz, 2022). Our observations of recruitment of DNMT3A to hypermethylated PMDs when H3K9me3 is reduced and H3K36me2 is elevated are consistent with these reports and also the observations that differences in the locations of PMDs between different head and neck tumours correlate with differences in H3K36me2 patterning (Zheng *et al*, 2023). They also suggest that DNMT3A re-localisation occurs following disruption of heterochromatin caused by DNMT1 loss. Indeed losses of H3K9me2/3 are previously reported in DNMT1 KO cells (Espada *et al*, 2004) and acute depletion or inhibition of DNMT1 reduces H3K9me3 levels and alters heterochromatin compartmentalisation (Scelfo *et al*, 2024; Spracklin *et al*, 2023). Further evidence of interplay between these two marks is provided by another recent study demonstrating that perturbation of H3K36me2 leads to alterations in H3K9me3 distribution (Padilla *et al*, 2024). It is interesting to note that in our study, DNMT3A ChIP-seq signal in hypermethylated PMDs is weaker than that observed for euchromatic domains. This suggests, that hypermethylated PMDs could retain some heterochromatic features that inhibit DNMT3A recruitment to some degree. We and others have reported that DNMT3A is also recruited to the polycomb-associated H2AK119ub mark through its N-terminal region (Chen *et al*, 2024; Gretarsson *et al*, 2024; Gu *et al*, 2022; Wapenaar *et al*, 2024; Weinberg *et al*, 2021). However, we do not observe the polycomb-associated H3K27me3 mark, which is generally tightly correlated with H2AK119ub (Ku *et al*, 2008), at hypermethylated PMDs suggesting that H2AK119ub does not play a role in the recruitment of DNMT3A to these regions.

We also note some degree of signal for both H3K4me3 and H3K36me3 within PMDs. Similar localisation of H3K4me3 to hypomethylated late replicating domains was previously reported (Du *et al*, 2021). However, the significance of these euchromatin-associated histone marks in heterochromatic domains remains unclear at present and will require further investigation.

DNA sequence has also been proposed to play a central role in the formation of PMDs. Within PMDs single CpG methylation levels correlate with CpG density (Gaidatzis *et al*, 2014) and hypomethylation is particularly apparent at CpGs that are flanked by A or T nucleotides and distant from other CpGs, termed solo-WCGW CpGs (Zhou *et al*., 2018). Methylation levels solo-WCpGWs negatively correlate with the number of mutations in tumours (Zhou *et al*., 2018) and also with population doublings *in vitro* (Endicott *et al*, 2022). A bias to loss of methylation to low CpG density regions is also reported for DNMT1 depletion by shRNA (Tiedemann *et al*., 2024). Here, we find that a PMD can convert to a non-PMD in association with alterations histone modifications and recruitment of *de novo* DNMTs without a change in sequence. This suggests that whereas sequence may play a role in PMD formation, it can be superseded by differences in chromatin structure and DNMT recruitment. The observation that tissue- and tumour subtype-specific PMDs exist (Salhab *et al*., 2018; Zheng *et al*., 2023) also demonstrates that the formation of PMDs is not solely driven by DNA sequence.

DNMT1 knockout HCT116 cells were generated by deletion of 3 exons of DNMT1 (Rhee *et al*., 2000). It was later shown that these cells express low levels of a truncated DNMT1 protein (Egger *et al*., 2006) and are hypomorphic. As with most mammalian cells, complete removal of DNMT1 is lethal to HCT116 cells (Chen *et al*, 2007). This means that it is possible that the formation of hypermethylated PMDs is a gain-of-function effect of the truncated DNMT1 allele. However, to our knowledge, the truncated protein has not been reported to present any additional gain-of-function effects.

In conclusion, we report an unexpected consequence of DNMT1 knockout in colorectal cancer cells is the gain of DNA methylation in several PMDs in association with the localisation of DNMT3A. These regions reveal a role of DNMT3A in maintaining DNA methylation homeostasis in cancer cells and, more generally, suggests that the lack of *de novo* activity in heterochromatin is a key factor in the formation of PMDs in cancer.

## Methods

### Cell culture

HCT116 and DNMT1 KO cells were gifts from B. Vogelstein (Rhee *et al*., 2000). Cells were cultured in McCoy’s 5A (Gibco) supplemented with 10% Fetal Calf Serum (Life Technologies) and penicillin–streptomycin antibiotics at 140 and 400 μg/ml, respectively at 37°C with 5% CO_2_. Cell lines were routinely tested for mycoplasma contamination. DNMT1 KO genotype was confirmed by western blot and sequencing.

### DNMT3A/B expression in cells

Stable cell lines ectopically expressing DNMT3A and DNMT3B were generated for HCT116 cells and DNMT1 KO cells. Cells were transfected via reverse transfection using FuGene HD transfection reagent (Promega) with PB-CAG-T7-DNMT3B2-IRES-puro or PB-CAG-T7-DNMT3A1-IRES-puro (Taglini *et al*., 2024) and a plasmid expressing piggyBac transposase. After transfection, cells were grown in media supplemented with 2μg/ml puromycin to generate stably expressing cell lines.

### DNA extraction

DNA was extracted from cell pellets snap-frozen in dry ice with ethanol. Cells pellets were resuspended in Genomic Lysis Buffer (300mM NaCl, 1% SDS, 20mM EDTA). The lysis mixture was syringed up and down using a 21G needle before being incubated at 55°C overnight with proteinase K. RNA was removed by incubation with RNase A/T1 Cocktail (Ambion) at 37 °C for 1 h, in between two phenol–chloroform extraction steps. DNA was quantified by Nanodrop 8000 spectrophotometer or Qubit fluorometer (Invitrogen) and purity was assessed using the Agilent 2100 BioAnalyzer.

### WGBS data generation

DNA (100ng) was bisulphite converted using the EZ DNA Methylation-Lightning kit, according to the manufacturer’s instructions. Libraries were generated using the TruSeq DNA Methylation kit, according to the manufacturer’s instructions. Sequencing was performed at the Edinburgh Clinical Research Facility. Libraries were sequenced using the Illumina NextSeq 500/550 High-Output v2 (150 cycles) (#FC-404-2002) on the NextSeq 550 platform (Illumina Inc, #SY-415-1002) over 1 flow cell. HCT116 cell WGBS used in this study have already were previously reported (Taglini *et al*., 2024) and DNMT1 KO cell data were generated in parallel.

### WGBS data processing

Fastq files were quality checked using FastQC (v0.11.4) and reads trimmed with TrimGalore (v0.4.1) using default settings. Reads were then mapped to a bisulfite converted genome (hg38) and duplicates were removed using bismark (v 0.18.1) with bowtie2 (v2.3.1) for paired end data (settings: -N0 -L20) (Krueger & Andrews, 2011; Langmead & Salzberg, 2012) before PCR duplicates were identified and removed using Bismark’s deduplicate_bismark command. Aligned BAM files were processed to report coverage and the number of methylated reads for each CpG observed. Forward and reverse strands were combined using Bismark’s methylation extractor and bismark2bedgraph modules with custom Python and AWK scripts. Processed WGBS files were assessed for conversion efficiency based on the proportion of methylated reads mapping to the phage-λ genome spike-in (>99.5% in all cases). For summary of WGBS alignment statistics, see *Supplementary Table 1*. Weighted mean methylation levels over genomic windows and domains were calculated by intersecting each window/domain with CpGs using bedtools intersect and groupby functions (v2.27.1).

### PMD annotation

To define PMDs, we used Methpipe (v5.0.0) (Decato et al., 2020) on CpG level WGBS data. Methpipe defined PMDs were further processed by trimming the domains obtained to the first and last CpG observed withing the domains, as well as by bridging nearby domains whose distance was smaller than 2x the window size used. The rest of the genome, not annotated as PMDs, was then assigned as highly methylated domains (HMDs) using bedtools (v2.27.1). We excluded poorly mapped regions of the genome using annotations of gaps and centromeres (using hg38 gap and centromere tracks downloaded from UCSC table browser). Regions annotated as heterochromatin, short arm and telomeres from the gaps track were merged with the centromeres track using bedtools merge with -d set to 10Mb. This merged file was then excluded from the PMD and HMD BED files using bedtools subtract. Annotated domains smaller than 200kb were also removed. After removal of the <200kb domains, the neighboring domains were then merged. Hypermethylated PMDs were defined as those ≥ 5% mean methylation in DNMT1 KO versus HCT116 cells.

### Pileup plots

ComputeMatrix from Deeptools (v3.5.0) was used to generate pileup plots and heatmaps. All domains were normalised to a size of 1Mb based on their start and end points using a window size of 10Kb (--binsize 10000 (-m 1000000). The flanks were plotted using the same window size and a maximum distance of 1Mb -b 1000000 -a 1000000).

### Repli-seq data generation

Repli-seq data was generated from cells at approximately 70% confluency to ensure most of the cells were actively cycling. To label DNA, cells were given fresh media supplemented with 100μM 5-Ethynyl-2’-deoxyuridine (EdU) and incubated at 37°C for 30 minutes. Cells were then collected by trypsinization along with media and a PBS wash of the flask to ensure non-adherent mitotic cells were not lost. The cell suspension was then centrifuged at 200g for 5 minutes and washed with 10ml PBS before being resuspended in 2.5ml PBS supplemented with 1% FCS. Cells were fixed by gently adding 7.5ml of 100% ethanol to a final concentration of 70% with gentle vortexing followed by incubation on ice for 30 mins and storage at –20°C.

Fixed cells were washed with 5ml ice cold PBS, and then resuspended in 5ml 100mM HCl and 0.015% pepsin (0.75mgr) in H_2_O before being incubated at 37°C for 30 minutes while rotating. Following centrifugation at 600g for 10 minutes, pellets were washed with 5 ml ice cold PBS and then resuspended in 1ml PBS along with 50μl Propidium Iodide (PI) and 20μl RNase A and incubated at room temperature for 30 minutes while protected from light. Extracted Nuclei were counted using a hematocytometer and suspended to approximately 3×10^6^ nuclei/ml to ensure optimal flow during FACS sorting. Nuclei were sorted on a FACSAria flow cytometer (BD Biosciences) in PBS supplemented with 0.25% BSA based on their DNA content using PI staining, into three fractions: Early, Middle and Late S-Phase (defined by equally splitting the window between the G1 and G2 peaks into three equal portions).

DNA was then extracted from sorted nuclei as described above except they were sonicated on a Bioruptor following proteinase K digestion (15 cycles of 30 seconds on – 30 seconds off, on high at 4°C). Precipitation of extracted DNA was then performed with linear acrylamide (LPA) carrier and isopropanol and resuspended in TE. DNA was then quantified using Qubit High Sensitivity reagents and sonicated on a Covaris E220 sonicator to yield 100-300bp fragments which were quality controlled on a Bioanalyser.

Re-precipitation of DNA was was then performed with LPA carrier and ethanol before being resuspended in water. A click reaction was then performed to conjugate biotin to the EdU by adding 3ul 2M TAB (0.4M), 30ul DMSO, 6ul 5mM Ascorbic Acid (2mM), 1.2ul 10mM Biotin Azide (0.8mM), 3ul 10mM CuTBTA (2mM) and 1.8ul H_2_O. Samples were then incubated overnight at room temperature while covered from light and cleaned up by ethanol precipitation with LPA carrier.

EdU containing DNA was then enriched using Myone C1 Streptavidin Dynabeads (Invitrogen) and a magnetic rack overnight at 4°C with rotation. Following this, the samples were washed at 4°C with: TSE-I (20 mM Tris-HCl, pH 8.1, 2 mM EDTA, 150 mM NaCl, 1% Triton, 0.1% (v/v) SDS) x2, TSE-II (20 mM Tris-HCl pH 8.1, 2 mM EDTA, 500 mM NaCl, 1% Triton, 0.1%(v/v) SDS) x1 and TE x1. Samples were eluted by adding water at 95°C and incubation with vortexing at 95°C for 10 mins. Successful enrichment of early and late replicating DNA was then validated by qPCR of early (BMP1, TEK, MELK, RTTN) and late replicating (NETO1, CDH8, DPPA2 and PTPRD) loci, based on a published protocol and HCT116 WT repli-seq data (Du *et al*., 2021; Marchal *et al*, 2018). Primer sequences used in this validation are included in *Supplementary Table 2*.

Repli-seq libraries were then prepapred using the Acel-NGS 1a Plus DNA library kit along with the 1A Plus Set A Indexing Kit following the manufacturer’s instruction (Swift Biosciences). However, as our DNA template contain Uracil due to the EdU incorporation, the kit’s polymerase, and subsequent buffer (Reagent W2, Buffer W3 and Enzyme W4) were replaced with KAPA HiFi HotStart Uracil+ Ready Mix Kit (Roche), which included an uracil tolerant polymerase. Libraries were assessed for size distribution on the Agilent Bioanalyser (Agilent Technologies) and quantified using the Qubit 2.0 Fluorometer. Repli-seq libraries were sent for whole genome sequencing at the Wellcome Trust Clinical Research Facility (WTCRF). The libraries were sequenced using the Illumina NextSeq 500/550 High-Output v2.5 (75 cycles) Kit on the Illumina NextSeq 550 platform over 2 flow cells to give 75bp single-end reads.

### Repli-seq data processing

Repli-seq fastq files were quality checked using FastQC (v0.11.4) and reads were trimmed using TrimGalore (v0.4.1) using default settings. Reads were then mapped to the genome (hg38) using bowtie2 for paired end data (version settings: -N1 -L 20 --no-unal -- no-mixed -- no-discordant -X 1000) (Langmead & Salzberg, 2012). Low mapping quality reads were removed using samtools (v1.6, settings: -bq10). Repli-seq data were analysed as previously described (Ryba *et al*, 2011). Briefly, read counts were generated for 10kb genomic windows using bedtools coverage (v2.27.1, settings: -counts) and normalised to reads per million (RPM). Replication timing was then calculated as the ratio between the early and late fraction RPM counts. Quantile normalisation of counts was then performed across samples and Loess smoothing was performed throughout all chromosomes using R (expect for the Y chromosome and mitochondrial DNA). The RPM counts over 10kb windows for early, middle, and late samples as well as the Loess smoothed replication timing counts were converted into BigWigs using bedGraphToBigWig from UCSC tools. For a summary of Repli-seq alignment statistics, see *Supplementary Table 3*.

### ChIP data generation

ChIP-seq was performed as previously described (Masalmeh *et al*., 2021; Taglini *et al*., 2024). For T7-DNMT3B and T7-DNMT3A ChIP-seq experiments, 1×10^7^ cells were harvested, washed and crosslinked with 1% methanol-free formaldehyde in PBS for 8 min at room temperature. Crosslinked cells were lysed for 10 min on ice in 50 μl of lysis buffer (50 mM Tris-HCl pH 8, 150 mM NaCl, 1 mM EDTA, 1% SDS) freshly supplemented with proteinase inhibitor (Sigma-Aldrich). IP dilution buffer (20 mM Tris-HCl pH 8, 150 mM NaCl, 1 mM EDTA, 0.1% Triton X-100) freshly supplemented with proteinase inhibitor, DTT and PMSF was added to the samples to reach a final volume of 500 μl. Chromatin was fragmented using Benzonase (Pchelintsev *et al*, 2016): samples were sonicated on ice with Soniprep 150 twice for 30 s to break up nuclei; then 200 U of Benzonase Nuclease (Sigma) and MgCl2 (final concentration 2.5 mM) were added and samples were incubated on ice for 15 min. The reaction was blocked by adding 10 μl of 0.5 M EDTA pH 8. Following centrifugation for 30 min at 18,407 g at 4 °C, supernatants were collected and supplemented with Triton X-100 (final concentration 1%) and 5% input aliquots were retained for later use. Protein A Dynabeads (Invitrogen) previously coupled with 10 μl of T7-Tag antibody per 1×10^7^ cells in blocking solution (1x PBS, 0.5% BSA) were added and the samples incubated overnight under rotation at 4 °C. Beads were then washed for 10 min at 4 °C with the following buffers: IP dilution buffer 1% Triton X-100 (20 mM Tris-HCl pH 8, 150 mM NaCl, 2 mM EDTA, 1% Triton X-100), buffer A (50mM HEPES pH 7.9, 500 mM NaCl, 1 mM EDTA, 1% Triton X-100, 0.1% Na-deoxycholate, 0.1% SDS), buffer B (20 mM Tris pH 8, 1 mM EDTA, 250 mM LiCl, 0.5% NP-40, 0.5% Na-deoxycholate), TE buffer (1mM EDTA pH 8, 10 mM Tris pH 8). Chromatin was eluted by incubating the beads in extraction buffer (0.1 M NaHCO3, 1% SDS) for 15 min at 37 °C. To reverse the cross-linking Tris-HCl pH 6.8 and NaCl were added to final concentrations of 130 mM and 300 mM respectively, and immunoprecipitations were incubated at 65 °C overnight. Samples were then incubated at 37 °C for 1 h after addition of 2 μl of RNase Cocktail Enzyme Mix (Ambion). Then 40 μg of Proteinase K (Roche) were added, followed by 2 h incubation at 55 °C. Input material was similarly de-crosslinked. Samples were purified with the MinElute PCR purification kit (QIAGEN). For ChIP-Rx-seq of ectopic T7-DNMT3B and T7-DNMT3A 20 μg of Spike-in chromatin (ActiveMotif 53083) was added to each sample after sonication. 2 μl of spike-in antibody per sample (ActiveMotif 61686) was also added in a ratio 1:5 versus the T7 antibody. A similar protocol was used for H3K4me3, H3K9me3, H3K27me3 and H3K36me3 ChIP-seq experiments, except: 0.5×10^7^ cells were harvested and crosslinked with 1% methanol-free formaldehyde in PBS for 5 min at room temperature. For H3K36me3 ChIP-Rx-seq crosslinked Drosophila S2 cells were spiked into samples before sonication at a ratio of 20:1 human to Drosophila cells. Following nuclei rupture by sonication on ice with Soniprep 150, chromatin was fragmented using Bioruptor Plus sonicator (Diagenode) for 40 cycles (30 s on/30 s off on high setting at 4 °C). 2 μl /1×10^6^ cells of the following antibodies were used for immunoprecipitations: H3K4me3 (EpiCypher 13-00041), H3K9me3 (Active Motif 39161), H3K27me3 (Cell Signaling Technology C36B11) and H3K36me3 (Abcam ab9050).

For H3K36me2, native ChIP-seq was performed adapting ultra-low-input (ULI) ChIP (Brind’Amour & Lorincz, 2022) as follows. Nuclei were isolated from 1×10^6^ unfixed cells using PBS 0.1% NP-40 and resuspended in 75 μl of Nuclear Isolation buffer (10mM Tris HCl, 0.1% NaDeOx, 0.1% Triton-100) and 75 μl of 2X MNase digestion Buffer (NEB). Chromatin was digested using MNase (NEB) for 6 min at 37 °C. Enzyme concentration was determined experimentally to obtain a majority of mononucleosomes and reaction stopped adding 5 μl of 0.5 M EDTA pH 8. Nuclei were lysed as follows: 30 μl of 5X Nuclear Lysis Buffer (5% NaDeOx, 5% Triton X-100) were added and samples vortexed, then 160 μl of ChIP Dilution buffer (0.1% Triton X-100, 20 mM Tris-HCl (pH 8.1), 150mM NaCl) were added and samples sonicated using Bioruptor Plus sonicator (Diagenode) for two cycles (30 sec on/30 sec off on high setting at 4°C). Following centrifugation for 30 min at maximum speed at 4 °C, supernatants were collected and 350 μl of ChIP Dilution buffer added. Half the sample was retained to check chromatin fragmentation and half processed for immunoprecipitation as previously described. 10% of input retained and antibody-bound-beads added for overnight incubation at 4 °C. 3 μl of anti-H3K36me2 (Invitrogen, T.571.7, MA5-14867) were used per sample. Washes and DNA elution were performed as described above.

Prior to library generation, qPCR was performed using SNAP-ChIP chromatin spike-in for specificity and enrichment of the antibodies for each sample as well as a positive and negative locus for each mark (H3K4me3, H3K27me3 and H3K9me3) based on ENCODE data for HCT116 cells (Consortium, 2012). Primers against MELK were used as a positive locus for H3K4me3 and as a negative locus for H3K9me3 and H3K27me3. Primers against ELAVL2 were used as a positive locus for H3K9me3 and as a negative locus for H3K4me3. Primers against MLLT3 were used as a positive locus for H3K27me3. Validation of successful DNMT-ChIP by qPCR was performed with a positive and negative locus for DNMT3A and DNMT3B (Masalmeh *et al*., 2021). Primers against DAZL were used as a positive locus for DNMT3A, and against TNFRSF1A for DNMT3B. Primers against BRCA2 were used as a negative locus for both DNMT3A and DNMT3B. Primer sequences used in this validation are included in *Supplementary Table 2*.

ChIP-seq libraries were prepared using the NEBNext Ultra II DNA Library Prep Kit for Illumina (NEB) and NEBNext Multiplex Oligos for Illumina (NEB) barcode adapters according to the manufacturer instructions. were used. Specifically, Illumina Index Primers Set 1 (E7335) for H3K36me3, and Unique Dual Index UMI Adaptors DNA Set 1 (E7395) for the other libraries. For histone modifications ChIP-seq, adapter-ligated DNA was size selected for an insert size of 150 bp using Agencourt AMPure XP beads. Libraries were quantified using the Qubit dsDNA HS or BR assay kit and assessed for size and quality using the Agilent Bioanalyser. H3K36me3 ChIP-Rx-seq libraries were sequenced using the NextSeq 500/550 high-output version 2.5 kit (75 bp single end reads). The other ChIP-seq libraries were sequenced using the NextSeq 2000 P3 (50 bp paired end reads). Libraries were combined into equimolar pools to run within individual flow cells. Sequencing was performed at the Edinburgh Clinical Research Facility. HCT116 ChIP-seq data used in this study have previously been reported (Masalmeh *et al*., 2021; Taglini *et al*., 2024), and DNMT1 KO data were generated in parallel.

### ChIP-seq data processing

Fastq files were quality checked using FastQC (v0.11.4) and reads trimmed using TrimGalore (v0.4.1). Reads were then mapped to the genome (hg38) using bowtie2 (v2.3.1 with settings: -N 1 -L 20 --no-unal) (Langmead & Salzberg, 2012) for paired end data. Low mapping quality reads or fragments were removed using samtools (v1.6 with settings -bq 10). Duplicate reads were removed using sambamba markdup (v0.5.9, for H3K4me3, H3K9me3, H3K27me3 and H3K36me3 ChIP-seq) or using UMI tools dedup function (v1.0.0 and setting: --paired, DNMT3A/B and H3K36me2 ChIP-seq). The number of reads pre- and post-alignment as well as the reads after deduplication and low mapping was then extracted and counted. Bed counts for 10 kb genomic windows and domains were generated from the bam files using bedtools coverage (seetings: *-counts*, v2.27.1). For paired-end data, the BAM file was first converted to a BED file of fragment locations using BEDtools bamtobed function. Coverage counts were scaled to counts per 10 million based on total number of mapped reads per sample and divided by the input read count to obtain a normalised read count. An offset of 0.5 was added to all windows prior to scaling and input normalisation to prevent intervals with zero reads in the input sample generating a normalised count of infinity. Regions where coverage was 0 in all samples were removed from the analysis. BigWig files for visualisation in the genome browser were generated, by calculating counts per million mapped reads using bamCoverage from deeptools (v3.2.0). Normalisation over inputs and means between replicas were then performed using bigwigCompare ratio and mean respectively. For a summary of ChIP-seq alignment statistics, see *Supplementary Tables 4-9*.

### Definition of H3K9me3-marked and H3K27me3-marked PMDs

Analysis of H3K9me3 and H3K27me3 enrichment across PMDs, showed a bimodal pattern for both marks, where high enrichment H3K9me3 PMDs showed low H3K27me3 enrichment and vice versa. K-means clustering was used to divide the PMDs into 2 clusters of H3K9me3-marked PMDs and H3K27me3-marked PMDs.

### Definition of H3K9me3 domains

H3K9me3 and H3K27me3 domains were called using a hidden Markov model as previously described (Spracklin & Pradhan, 2020). Normalised mean H3K9me3 ChIP-seq coverage in 25 kb genomic windows was analysed using the bigwig_hmm.py script (https://github.com/gspracklin/hmm_bigwigs) to define two states (-n 2). We excluded poorly mapped regions of the genome from these domains using annotations of gaps and centromeres from the UCSC browser (hg38 gap and centromere tracks). Annotations were downloaded from the UCSC table browser. Regions annotated as heterochromatin, short arm and telomeres from the gaps track were merged with the centromeres track using BEDtools merge with -d set to 10 Mb. This merged file was then excluded from the domains BED files using BEDtools subtract.

### RNA-seq data generation

RNA extraction was performed from snap-frozen cell pellets using RNeasy Plus Mini Kit (QIAGEN) following the manufacturer’s instructions. Quantity and quality of RNA samples were assessed using the Nanodrop spectrophotometer (Nanodrop ND-1000, Thermo Scientific) and Qubit fluorometer (Invitrogen). Size distribution of RNA fragments and the RNA integrity number (RIN) value were determined by using Agilent 2100 BioAnalyzer with the RNA nano chip. RNA samples for whole genome sequencing were sent to Edinburgh Genomics for library preparation and sequence data generation. Library preparation was performed using the TruSeq stranded mRNA-seq library kit. The libraries were then sequenced on NovaSeq with 50bp paired end (PE) reads and aiming for 50M reads per sample.

### RNA-seq data processing and analysis

RNA-seq fastq files were quality checked using FastQC (v0.11.4) and reads were trimmed using TrimGalore (v0.4.1) and default settings. Reads were then mapped to the genome (hg38) using bowtie2 for paired end data (2.3.1, settings: -N1 -L 20 --no-unal -- no-mixed -- no-discordant -X 1000) (Langmead & Salzberg, 2012). Low mapping quality reads were removed using samtools (settings: -bq10). The bam file was then converted into bed using bedtools and CPM values calculated. BigWig files were then generated using the ucsc toolset, for visualisation in the genome browser. FeatureCounts (settings: -T 4 -t exon -p, from subread package) was then used to count how many reads align to each gene. For a summary of RNA-seq alignment statistics, see *Supplementary Table 10*. Part of the HCT116 RNA-seq data used here was previously reported (Taglini *et al*., 2024) and the rest of the data was generated in parallel.

Statistical analysis to detect differentially expressed genes between genotypes was performed using the edgeR package in R (Robinson *et al*, 2010). Normalisation factors were then calculated using the trimmed mean of M values (TMM) normalisation method and differential expression analysis was performed using a general linear model (GLM). Statistically significant differentially expressed genes were defined as the genes that demonstrated a false discovery rate (FDR) < 0.05 and an absolute log fold change > 1.

### Annotation of functional elements

Gene positions in genome browser plots are hg38 RefSeq genes taken from UCSC. CpG island positions were taken from Illingworth *et al*, converted to hg38 using the UCSC liftover tool (Illingworth *et al*, 2010).

## Statistical analysis

Statistical tests were performed in R (3.3.3) or GraphPad Prism (9.0.0).

## Code availability

Custom scripts used in the analysis of data are available from the authors upon request.

## Data availability

All sequencing data that were generated during this study will be deposited in GEO and made available upon publication.

## Acknowledgements

Edinburgh Clinical Research Facility Genetics Core and Edinburgh Genomics for conducting the high-throughput sequencing used in this study. This work has made use of the resources provided by the University of Edinburgh digital research services and the MRC IGC compute cluster. DS is a Cancer Research UK Career Development fellow (reference C47648/A20837), and work in his laboratory is also supported by an MRC university grant to the MRC Human Genetics Unit. IK was funded by a studentship from Cancer Research UK (C157/A25186) as well as the AG Leventis Foundation (18736).

## Contributions

Ioannis Kafetzopoulos: Conceptualisation; Formal analysis; Investigation; Writing—original draft; Writing—review and editing. Francesca Taglini: Conceptualisation; Formal analysis; Investigation; Writing—review and editing. Hazel Davidson-Smith: Investigation. Duncan Sproul: Conceptualisation; Formal analysis; Supervision; Funding acquisition; Writing— original draft; Writing—review and editing.

## Competing Interests

The authors declare no competing interests.

## Notes

### Competing Interest Statement

The authors have declared no competing interest.

## References

Aran D, Toperoff G, Rosenberg M, Hellman A (2011) Replication timing-related and gene body-specific methylation of active human genes. Hum Mol Genet 20: 670–680

Baubec T, Colombo DF, Wirbelauer C, Schmidt J, Burger L, Krebs AR, Akalin A, Schubeler D (2015) Genomic profiling of DNA methyltransferases reveals a role for DNMT3B in genic methylation. Nature 520: 243–247

Berman BP, Weisenberger DJ, Aman JF, Hinoue T, Ramjan Z, Liu Y, Noushmehr H, Lange CP, van Dijk CM, Tollenaar RA et al (2011) Regions of focal DNA hypermethylation and long-range hypomethylation in colorectal cancer coincide with nuclear lamina-associated domains. Nat Genet 44: 40–46

Besselink N, Keijer J, Vermeulen C, Boymans S, de Ridder J, van Hoeck A, Cuppen E, Kuijk E (2023) The genome-wide mutational consequences of DNA hypomethylation. Sci Rep 13: 6874

Brind’Amour J, Lorincz MC (2022) Profiling Histone Methylation in Low Numbers of Cells. Methods Mol Biol 2529: 229–251

Charlton J, Downing TL, Smith ZD, Gu H, Clement K, Pop R, Akopian V, Klages S, Santos DP, Tsankov AM et al (2018) Global delay in nascent strand DNA methylation. Nat Struct Mol Biol 25: 327–332

Chen T, Hevi S, Gay F, Tsujimoto N, He T, Zhang B, Ueda Y, Li E (2007) Complete inactivation of DNMT1 leads to mitotic catastrophe in human cancer cells. Nat Genet 39: 391–396

Chen X, Guo Y, Zhao T, Lu J, Fang J, Wang Y, Wang GG, Song J (2024) Structural basis for the H2AK119ub1-specific DNMT3A-nucleosome interaction. Nat Commun 15: 6217

Consortium EP (2012) An integrated encyclopedia of DNA elements in the human genome. Nature 489: 57–74

Cruickshank MN, Oshlack A, Theda C, Davis PG, Martino D, Sheehan P, Dai Y, Saffery R, Doyle LW, Craig JM (2013) Analysis of epigenetic changes in survivors of preterm birth reveals the effect of gestational age and evidence for a long term legacy. Genome Med 5: 96

Dahlet T, Argueso Lleida A, Al Adhami H, Dumas M, Bender A, Ngondo RP, Tanguy M, Vallet J, Auclair G, Bardet AF et al (2020) Genome-wide analysis in the mouse embryo reveals the importance of DNA methylation for transcription integrity. Nat Commun 11: 3153

Decato BE, Qu J, Ji X, Wagenblast E, Knott SRV, Hannon GJ, Smith AD (2020) Characterization of universal features of partially methylated domains across tissues and species. Epigenetics Chromatin 13: 39

Du Q, Bert SA, Armstrong NJ, Caldon CE, Song JZ, Nair SS, Gould CM, Luu PL, Peters T, Khoury A et al (2019) Replication timing and epigenome remodelling are associated with the nature of chromosomal rearrangements in cancer. Nat Commun 10: 416

Du Q, Smith GC, Luu PL, Ferguson JM, Armstrong NJ, Caldon CE, Campbell EM, Nair SS, Zotenko E, Gould CM et al (2021) DNA methylation is required to maintain both DNA replication timing precision and 3D genome organization integrity. Cell Rep 36: 109722

Egger G, Jeong S, Escobar SG, Cortez CC, Li TW, Saito Y, Yoo CB, Jones PA, Liang G (2006) Identification of DNMT1 (DNA methyltransferase 1) hypomorphs in somatic knockouts suggests an essential role for DNMT1 in cell survival. Proc Natl Acad Sci U S A 103: 14080–14085

Endicott JL, Nolte PA, Shen H, Laird PW (2022) Cell division drives DNA methylation loss in late-replicating domains in primary human cells. Nat Commun 13: 6659

Espada J, Ballestar E, Fraga MF, Villar-Garea A, Juarranz A, Stockert JC, Robertson KD, Fuks F, Esteller M (2004) Human DNA methyltransferase 1 is required for maintenance of the histone H3 modification pattern. J Biol Chem 279: 37175–37184

Feinberg AP, Vogelstein B (1983) Hypomethylation distinguishes genes of some human cancers from their normal counterparts. Nature 301: 89–92

Gaidatzis D, Burger L, Murr R, Lerch A, Dessus-Babus S, Schubeler D, Stadler MB (2014) DNA sequence explains seemingly disordered methylation levels in partially methylated domains of Mammalian genomes. PLoS Genet 10: e1004143

Gama-Sosa MA, Slagel VA, Trewyn RW, Oxenhandler R, Kuo KC, Gehrke CW, Ehrlich M (1983) The 5-methylcytosine content of DNA from human tumors. Nucleic Acids Res 11: 6883–6894

Gretarsson KH, Abini-Agbomson S, Gloor SL, Weinberg DN, McCuiston JL, Kumary VUS, Hickman AR, Sahu V, Lee R, Xu X et al (2024) Cancer-associated DNA hypermethylation of Polycomb targets requires DNMT3A dual recognition of histone H2AK119 ubiquitination and the nucleosome acidic patch. Sci Adv 10: eadp0975

Gu T, Hao D, Woo J, Huang TW, Guo L, Lin X, Guzman AG, Tovy A, Rosas C, Jeong M et al (2022) The disordered N-terminal domain of DNMT3A recognizes H2AK119ub and is required for postnatal development. Nat Genet 54: 625–636

Hon GC, Hawkins RD, Caballero OL, Lo C, Lister R, Pelizzola M, Valsesia A, Ye Z, Kuan S, Edsall LE et al (2012) Global DNA hypomethylation coupled to repressive chromatin domain formation and gene silencing in breast cancer. Genome Res 22: 246–258

Hovestadt V, Jones DT, Picelli S, Wang W, Kool M, Northcott PA, Sultan M, Stachurski K, Ryzhova M, Warnatz HJ et al (2014) Decoding the regulatory landscape of medulloblastoma using DNA methylation sequencing. Nature 510: 537–541

Howard G, Eiges R, Gaudet F, Jaenisch R, Eden A (2008) Activation and transposition of endogenous retroviral elements in hypomethylation induced tumors in mice. Oncogene 27: 404–408

Illingworth RS, Gruenewald-Schneider U, Webb S, Kerr AR, James KD, Turner DJ, Smith C, Harrison DJ, Andrews R, Bird AP (2010) Orphan CpG islands identify numerous conserved promoters in the mammalian genome. PLoS Genet 6: e1001134

Janssen SM, Lorincz MC (2022) Interplay between chromatin marks in development and disease. Nat Rev Genet 23: 137–153

Jones PA, Liang G (2009) Rethinking how DNA methylation patterns are maintained. Nat Rev Genet 10: 805–811

Krueger F, Andrews SR (2011) Bismark: a flexible aligner and methylation caller for Bisulfite-Seq applications. Bioinformatics 27: 1571–1572

Ku M, Koche RP, Rheinbay E, Mendenhall EM, Endoh M, Mikkelsen TS, Presser A, Nusbaum C, Xie X, Chi AS et al (2008) Genomewide analysis of PRC1 and PRC2 occupancy identifies two classes of bivalent domains. PLoS Genet 4: e1000242

Langmead B, Salzberg SL (2012) Fast gapped-read alignment with Bowtie 2. Nat Methods 9: 357–359

Li E, Bestor TH, Jaenisch R (1992) Targeted mutation of the DNA methyltransferase gene results in embryonic lethality. Cell 69: 915–926

Lister R, Pelizzola M, Dowen RH, Hawkins RD, Hon G, Tonti-Filippini J, Nery JR, Lee L, Ye Z, Ngo QM et al (2009) Human DNA methylomes at base resolution show widespread epigenomic differences. Nature 462: 315–322

Lister R, Pelizzola M, Kida YS, Hawkins RD, Nery JR, Hon G, Antosiewicz-Bourget J, O’Malley R, Castanon R, Klugman S et al (2011) Hotspots of aberrant epigenomic reprogramming in human induced pluripotent stem cells. Nature 471: 68–73

Marchal C, Sasaki T, Vera D, Wilson K, Sima J, Rivera-Mulia JC, Trevilla-Garcia C, Nogues C, Nafie E, Gilbert DM (2018) Genome-wide analysis of replication timing by next-generation sequencing with E/L Repli-seq. Nat Protoc 13: 819–839

Masalmeh RHA, Taglini F, Rubio-Ramon C, Musialik KI, Higham J, Davidson-Smith H, Kafetzopoulos I, Pawlicka KP, Finan HM, Clark R et al (2021) De novo DNA methyltransferase activity in colorectal cancer is directed towards H3K36me3 marked CpG islands. Nat Commun 12: 694

Ming X, Zhang Z, Zou Z, Lv C, Dong Q, He Q, Yi Y, Li Y, Wang H, Zhu B (2020) Kinetics and mechanisms of mitotic inheritance of DNA methylation and their roles in aging-associated methylome deterioration. Cell Res 30: 980–996

Nicetto D, Zaret KS (2019) Role of H3K9me3 heterochromatin in cell identity establishment and maintenance. Curr Opin Genet Dev 55: 1–10

Okano M, Bell DW, Haber DA, Li E (1999) DNA methyltransferases Dnmt3a and Dnmt3b are essential for de novo methylation and mammalian development. Cell 99: 247–257

Padilla R, Shipman GA, Horth C, Bareke E, Majewski J (2024) H3K36 Methylation - a Guardian of Epigenome Integrity. bioRxiv: 2024.2008.2010.607446

Pchelintsev NA, Adams PD, Nelson DM (2016) Critical Parameters for Efficient Sonication and Improved Chromatin Immunoprecipitation of High Molecular Weight Proteins. PLoS One 11: e0148023

Petryk N, Bultmann S, Bartke T, Defossez PA (2021) Staying true to yourself: mechanisms of DNA methylation maintenance in mammals. Nucleic Acids Res 49: 3020–3032

Raddatz G, Gao Q, Bender S, Jaenisch R, Lyko F (2012) Dnmt3a protects active chromosome domains against cancer-associated hypomethylation. PLoS Genet 8: e1003146

Rhee I, Jair KW, Yen RW, Lengauer C, Herman JG, Kinzler KW, Vogelstein B, Baylin SB, Schuebel KE (2000) CpG methylation is maintained in human cancer cells lacking DNMT1. Nature 404: 1003–1007

Robinson MD, McCarthy DJ, Smyth GK (2010) edgeR: a Bioconductor package for differential expression analysis of digital gene expression data. Bioinformatics 26: 139–140

Ryba T, Battaglia D, Pope BD, Hiratani I, Gilbert DM (2011) Genome-scale analysis of replication timing: from bench to bioinformatics. Nat Protoc 6: 870–895

Salhab A, Nordstrom K, Gasparoni G, Kattler K, Ebert P, Ramirez F, Arrigoni L, Muller F, Polansky JK, Cadenas C et al (2018) A comprehensive analysis of 195 DNA methylomes reveals shared and cell-specific features of partially methylated domains. Genome Biol 19: 150

Scelfo A, Barra V, Abdennur N, Spracklin G, Busato F, Salinas-Luypaert C, Bonaiti E, Velasco G, Bonhomme F, Chipont A et al (2024) Tunable DNMT1 degradation reveals DNMT1/DNMT3B synergy in DNA methylation and genome organization. J Cell Biol 223

Shirane K, Miura F, Ito T, Lorincz MC (2020) NSD1-deposited H3K36me2 directs de novo methylation in the mouse male germline and counteracts Polycomb-associated silencing. Nat Genet 52: 1088–1098

Spada F, Haemmer A, Kuch D, Rothbauer U, Schermelleh L, Kremmer E, Carell T, Langst G, Leonhardt H (2007) DNMT1 but not its interaction with the replication machinery is required for maintenance of DNA methylation in human cells. J Cell Biol 176: 565–571

Spracklin G, Abdennur N, Imakaev M, Chowdhury N, Pradhan S, Mirny LA, Dekker J (2023) Diverse silent chromatin states modulate genome compartmentalization and loop extrusion barriers. Nat Struct Mol Biol 30: 38–51

Spracklin G, Pradhan S (2020) Protect-seq: genome-wide profiling of nuclease inaccessible domains reveals physical properties of chromatin. Nucleic Acids Res 48: e16

Stewart-Morgan KR, Requena CE, Flury V, Du Q, Heckhausen Z, Hajkova P, Groth A (2023) Quantifying propagation of DNA methylation and hydroxymethylation with iDEMS. Nat Cell Biol 25: 183–193

Stroud H, Su SC, Hrvatin S, Greben AW, Renthal W, Boxer LD, Nagy MA, Hochbaum DR, Kinde B, Gabel HW et al (2017) Early-Life Gene Expression in Neurons Modulates Lasting Epigenetic States. Cell 171: 1151–1164 e1116

Suzuki MM, Bird A (2008) DNA methylation landscapes: provocative insights from epigenomics. Nat Rev Genet 9: 465–476

Taglini F, Kafetzopoulos I, Rolls W, Musialik KI, Lee HY, Zhang Y, Marenda M, Kerr L, Finan H, Rubio-Ramon C et al (2024) DNMT3B PWWP mutations cause hypermethylation of heterochromatin. EMBO Rep 25: 1130–1155

Tibben BM, Rothbart SB (2024) Mechanisms of DNA Methylation Regulatory Function and Crosstalk with Histone Lysine Methylation. J Mol Biol 436: 168394

Tiedemann RL, Hrit J, Du Q, Wiseman AK, Eden HE, Dickson BM, Kong X, Chomiak AA, Vaughan RM, Tibben BM et al (2024) UHRF1 ubiquitin ligase activity supports the maintenance of low-density CpG methylation. Nucleic Acids Res

Wapenaar H, Clifford G, Rolls W, Pasquier M, Burdett H, Zhang Y, Deak G, Zou J, Spanos C, Taylor MRD et al (2024) The N-terminal region of DNMT3A engages the nucleosome surface to aid chromatin recruitment. EMBO Rep

Weinberg DN, Papillon-Cavanagh S, Chen H, Yue Y, Chen X, Rajagopalan KN, Horth C, McGuire JT, Xu X, Nikbakht H et al (2019) The histone mark H3K36me2 recruits DNMT3A and shapes the intergenic DNA methylation landscape. Nature 573: 281–286

Weinberg DN, Rosenbaum P, Chen X, Barrows D, Horth C, Marunde MR, Popova IK, Gillespie ZB, Keogh MC, Lu C et al (2021) Two competing mechanisms of DNMT3A recruitment regulate the dynamics of de novo DNA methylation at PRC1-targeted CpG islands. Nat Genet 53: 794–800

Wild L, Flanagan JM (2010) Genome-wide hypomethylation in cancer may be a passive consequence of transformation. Biochim Biophys Acta 1806: 50–57

Yamanaka S, Nishihara H, Toh H, Eijy Nagai LA, Hashimoto K, Park SJ, Shibuya A, Suzuki AM, Tanaka Y, Nakai K et al (2019) Broad Heterochromatic Domains Open in Gonocyte Development Prior to De Novo DNA Methylation. Dev Cell 51: 21–34 e25

Zheng Y, Ziman B, Ho AS, Sinha UK, Xu LY, Li EM, Koeffler HP, Berman BP, Lin DC (2023) Comprehensive analyses of partially methylated domains and differentially methylated regions in esophageal cancer reveal both cell-type- and cancer-specific epigenetic regulation. Genome Biol 24: 193

Zhou W, Dinh HQ, Ramjan Z, Weisenberger DJ, Nicolet CM, Shen H, Laird PW, Berman BP (2018) DNA methylation loss in late-replicating domains is linked to mitotic cell division. Nat Genet 50: 591–602

