## Supplementary Figures and Legends for "DNMT1 loss leads to hypermethylation of a subset of late replicating domains by DNMT3A"

**Kafetzopoulos et al., supplementary materials**

**Contents**

*Figure S1* *page 2*
*Figure S2* *page 4*
*Figure S3* *page 6*
*Figure S4* *page 8*
*Figure S5* *page 10*
*Supplementary tables 1 to 10 are provided as an excel file.*

Fig. S1

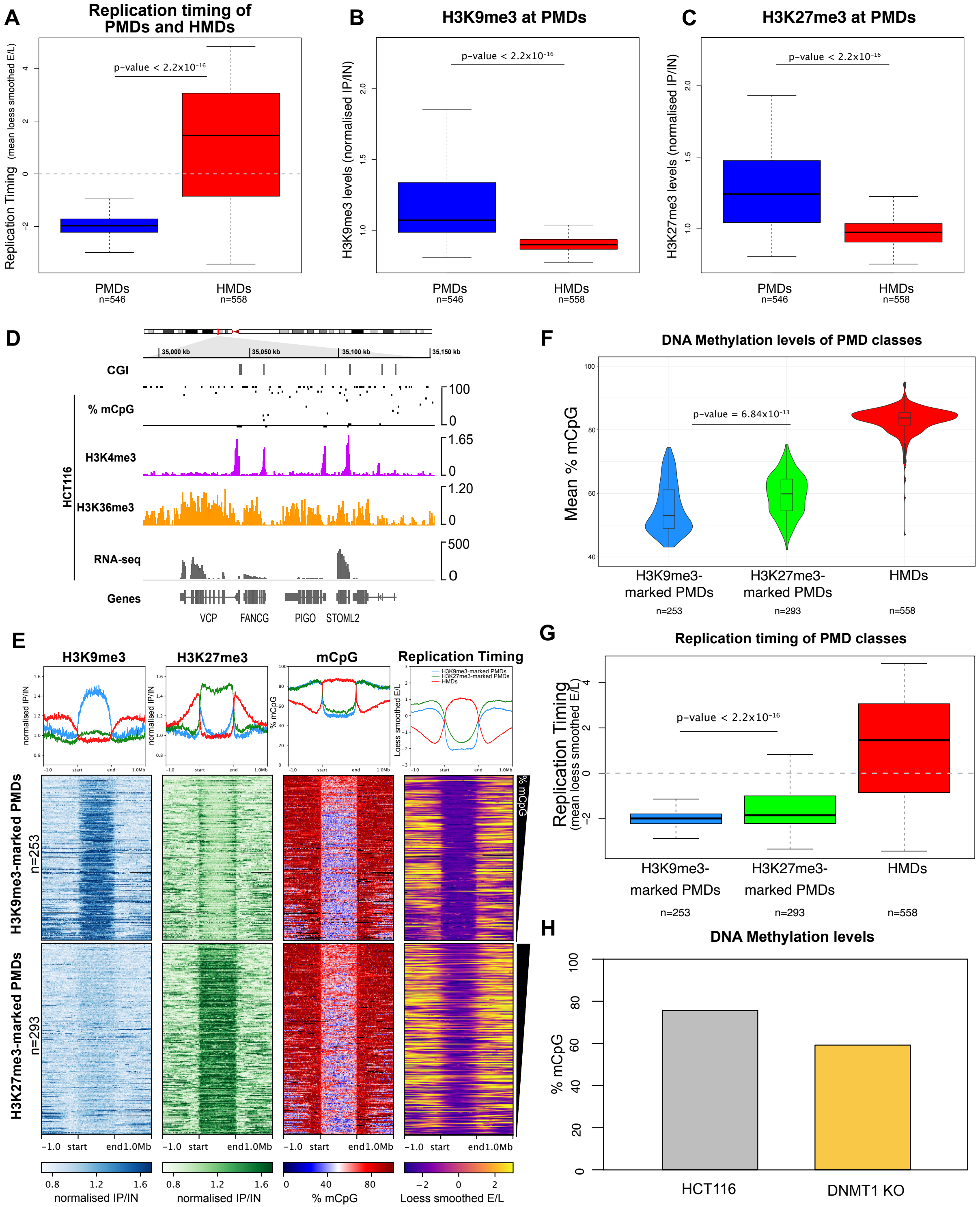
A
Replication timing of PMDs and HMDs
Replication Timing (mean loess smoothed E/L)
p-value < 2.2x10-16
PMDs
n=546
HMDs
n=558
B
H3K9me3 at PMDs
H3K9me3 levels (normalised IP/IN)
p-value < 2.2x10-16
PMDs
n=546
HMDs
n=558
C
H3K27me3 at PMDs
H3K27me3 levels (normalised IP/IN)
p-value < 2.2x10-16
PMDs
n=546
HMDs
n=558
D
35,000 kb
35,050 kb
35,100 kb
35,150 kb
CGI
% mCpG
H3K4me3
H3K36me3
RNA-seq
Genes
HCT116
VCP
FANCG
PIGO
STOML2
E
H3K9me3
H3K27me3
mCpG
Replication Timing
H3K9me3-marked PMDs
H3K27me3-marked PMDs
HMDs
H3K9me3-marked PMDs n=253
H3K27me3-marked PMDs n=293
normalised IP/IN
% mCpG
Loess smoothed E/L
-1.0 start end 1.0Mb
F
DNA Methylation levels of PMD classes
Mean % mCpG
p-value = 6.84x10-13
H3K9me3-marked PMDs
H3K27me3-marked PMDs
HMDs
n=253
n=293
n=558
G
Replication timing of PMD classes
Replication Timing (mean loess smoothed E/L)
p-value < 2.2x10-16
H3K9me3-marked PMDs
H3K27me3-marked PMDs
HMDs
n=253
n=293
n=558
H
DNA Methylation levels
% mCpG
HCT116
DNMT1 KO

**Figure S1. Ablation of DNMT1 leads to preferential hypomethylation of partially methylated domains.**

(**A**) Boxplot showing replication timing of HCT116 PMDs (n = 546 domains) and HMDs (n = 558 domains). Replication timing data are mean loess smoothed repli-seq early/late ratios over 10 kb. (**B**) Boxplot showing H3K9me3 levels of HCT116 PMDs (n = 546 domains) and HMDs (n = 558 domains). ChIP-seq data are mean normalised IP/IN. (**C**) Boxplot showing H3K27me3 levels of HCT116 PMDs (n = 546 domains) and HMDs (n = 558 domains). ChIP-seq data are mean normalised IP/IN. (**D**) Representative genomic locus showing histone marks at genes in HCT116 cells. Genome browser plot showing DNA methylation levels (mC) alongside HCT116 histone modification ChIP-seq and gene expression. DNA methylation levels are plotted for individual CpGs with coverage $\geq 5$. ChIP-seq are normalised $\log_{10}$ IP/IN. RNA-seq are mean logRPKM. CGI = CpG islands. (**E**) Heatmaps and pileup plots of HCT116 H3K9me3, H3K27me3 and DNA methylation levels alongside replication timing for H3K9me3-marked PMDs (n= 253) and H3K27me3-marked PMDs (n= 293). ChIP-seq data are mean normalised IP/IN, DNA methylation levels are mean % mCpG over 10kb. Replication timing data are mean loess smoothed repli-seq early/late ratios over 10kb. PMDs are aligned and scaled to the start and end points of each domain and ranked based on their mean methylation levels in HCT116 cells. (**F**) Violin plot showing mean HCT116 DNA methylation levels at H3K9me3 PMDs (n = 253 domains), H3K27me3 PMDs (n = 293 domains) and HMDs (n = 558 domains). (**G**) Boxplot showing replication timing of HCT116 H3K9me3 PMDs (n = 253 domains), H3K27me3 PMDs (n = 293 domains) and HMDs (n = 558 domains). Replication timing data are mean loess smoothed repli-seq early/late ratios over 10 kb. (**H**) Total DNA methylation levels are reduced in DNMT1 KO cells. Barplot of total methylated CpG levels estimated by WGBS. For boxplots: Lines = median; box = 25th–75th percentile; whiskers = 1.5 × interquartile range from box. All p-values are from two-sided Wilcoxon rank sum tests. All histone ChIP-seq and repli-seq data shown are derived from the mean of two biological replicates.

Fig. S2

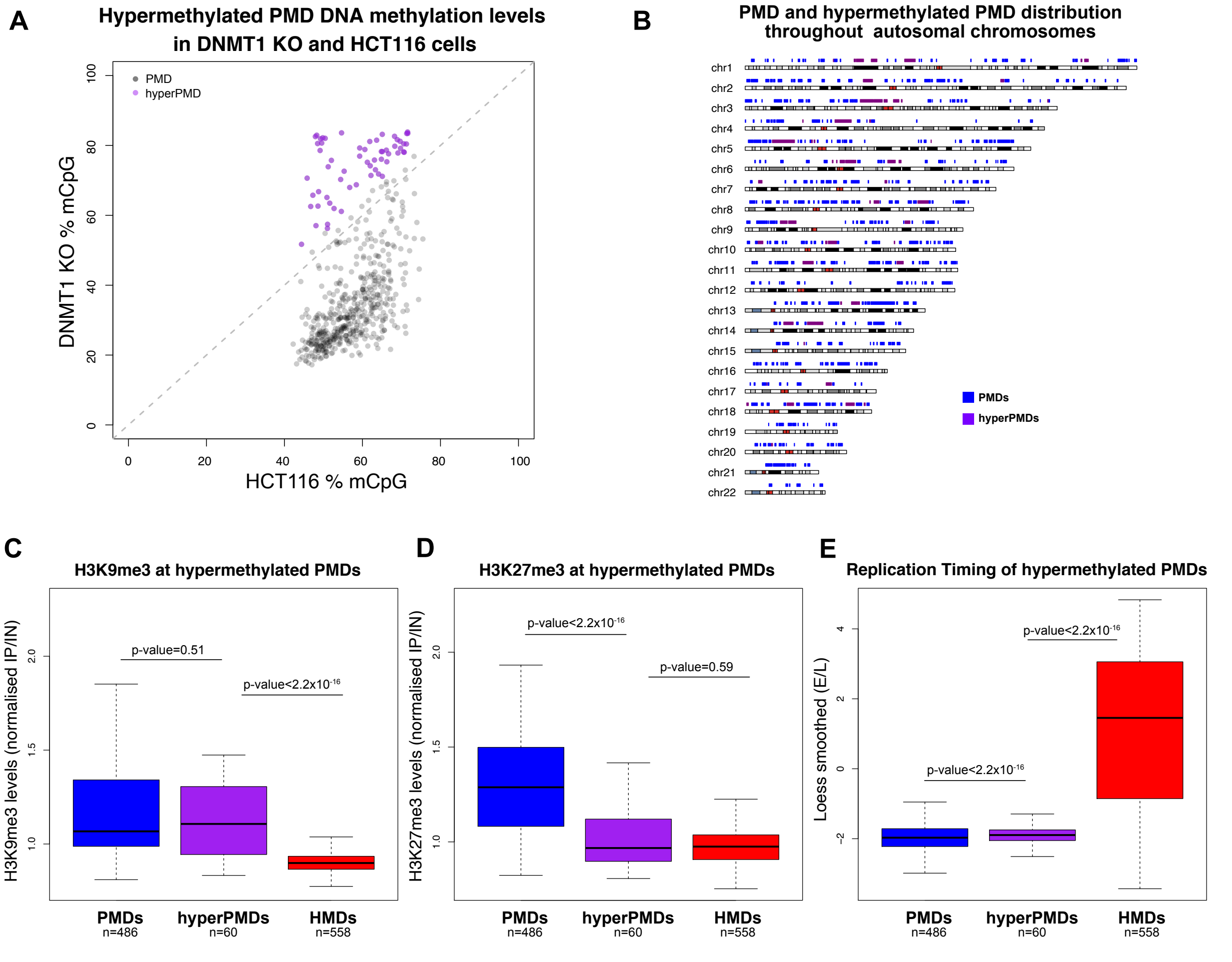
A
Hypermethylated PMD DNA methylation levels in DNMT1 KO and HCT116 cells
PMD
hyperPMD
DNMT1 KO % mCpG
HCT116 % mCpG
B
PMD and hypermethylated PMD distribution throughout autosomal chromosomes
chr1
chr2
chr3
chr4
chr5
chr6
chr7
chr8
chr9
chr10
chr11
chr12
chr13
chr14
chr15
chr16
chr17
chr18
chr19
chr20
chr21
chr22
PMDs
hyperPMDs
C
H3K9me3 at hypermethylated PMDs
H3K9me3 levels (normalised IP/IN)
p-value=0.51
p-value<2.2x10-16
PMDs n=486
hyperPMDs n=60
HMDs n=558
D
H3K27me3 at hypermethylated PMDs
H3K27me3 levels (normalised IP/IN)
p-value<2.2x10-16
p-value=0.59
PMDs n=486
hyperPMDs n=60
HMDs n=558
E
Replication Timing of hypermethylated PMDs
Loess smoothed (E/L)
p-value<2.2x10-16
p-value<2.2x10-16
PMDs n=486
hyperPMDs n=60
HMDs n=558

**Figure S2. A subset of H3K9me3-marked PMDs are hypermethylated in DNMT1 knockout cells.**

(**A**) Scatter plot of mean methylation levels at PMDs in DNMT1 KO cells versus HCT116 cells highlighting hypermethylated PMDs (hyperPMDs). (**B**) Ideogram showing genomic distribution and size of hypermethylated PMDs. (**C**) Boxplot showing HCT116 H3K9me3 levels at hypermethylated PMDs (n = 60 Domains compared to other PMDs (n = 486 domains) and HMDs (n = 558 domains). ChIP-seq data are mean normalised IP/IN. (**D**) Boxplot showing HCT116 H3K27me3 levels at hypermethylated PMDs (n = 60 Domains compared to other PMDs (n = 486 domains) and HMDs (n = 558 domains). ChIP-seq data are mean normalised IP/IN. (**E**) Boxplot showing HCT116 replication timing at hypermethylated PMDs (n = 60 Domains compared to other PMDs (n = 486 domains) and HMDs (n = 558 domains). Replication timing data are mean loess smoothed repli-seq early/late ratios over 10kb. For boxplots: Lines = median; box = 25th–75th percentile; whiskers = 1.5 × interquartile range from box. All p-values are from two-sided Wilcoxon rank sum tests. All histone ChIP-seq and repli-seq data shown are derived from the mean of two biological replicates.

### Fig. S3

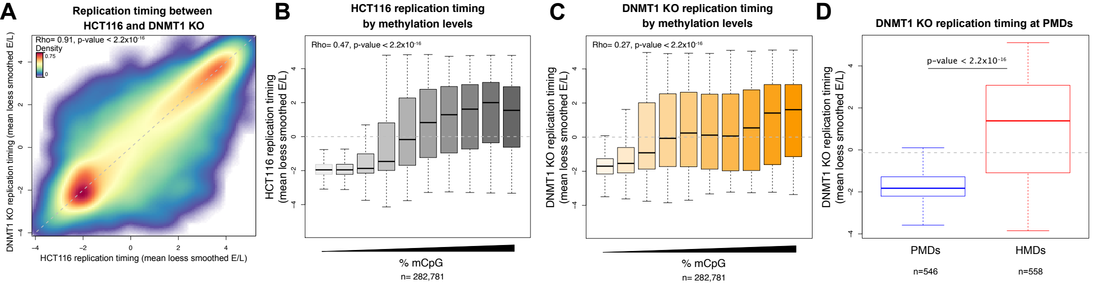
A
Replication timing between HCT116 and DNMT1 KO
Rho= 0.91, p-value < 2.2x10-16
Density
0.75
0
DNMT1 KO replication timing (mean loess smoothed E/L)
HCT116 replication timing (mean loess smoothed E/L)
B
HCT116 replication timing by methylation levels
Rho= 0.47, p-value < 2.2x10-16
HCT116 replication timing (mean loess smoothed E/L)
% mCpG
n= 282,781
C
DNMT1 KO replication timing by methylation levels
Rho= 0.27, p-value < 2.2x10-16
DNMT1 KO replication timing (mean loess smoothed E/L)
% mCpG
n= 282,781
D
DNMT1 KO replication timing at PMDs
p-value < 2.2x10-16
DNMT1 KO replication timing (mean loess smoothed E/L)
PMDs
HMDs
n=546
n=558

**Figure S3. The replication timing of hypermethylated PMDs remains similar in DNMT1 KO cells.**

(**A**) Density scatter plot showing genome-wide correlation between replication timing in DNMT1 KO and HCT116 cells. Replication timing data are mean loess smoothed repli-seq early/late ratios in 10kb windows. Spearman's correlation (Rho) and associated p-value is shown. (**B**) Boxplot showing HCT116 replication timing in 10 kb genomic windows divided in deciles according to their mean DNA methylation levels in HCT116. Replication timing data are mean loess smoothed repli-seq early/late ratios over 10 kb. Spearman's correlation coefficient (Rho) is shown alongside its associated p-value and n is the number of windows analysed. (**C**) Boxplot showing DNMT1 KO replication timing in 10 kb genomic windows divided in deciles according to their mean DNA methylation levels in DNMT1 KO. Replication timing data are mean loess smoothed repli-seq early/late ratios over 10 kb. Spearman's correlation coefficient (Rho) is shown alongside its associated p-value and n is the number of windows analysed. (**D**) Boxplot showing replication timing of HCT116 PMDs (n = 486 domains) and HMDs (n = 558 domains). Replication timing data are mean loess smoothed repli-seq early/late ratios over 10 kb. For boxplots: Lines = median; box = 25th–75th percentile; whiskers = 1.5 × interquartile range from box. All p-values are from two-sided Wilcoxon rank sum tests. All repli-seq data shown are derived from the mean of two biological replicates.

Fig. S4

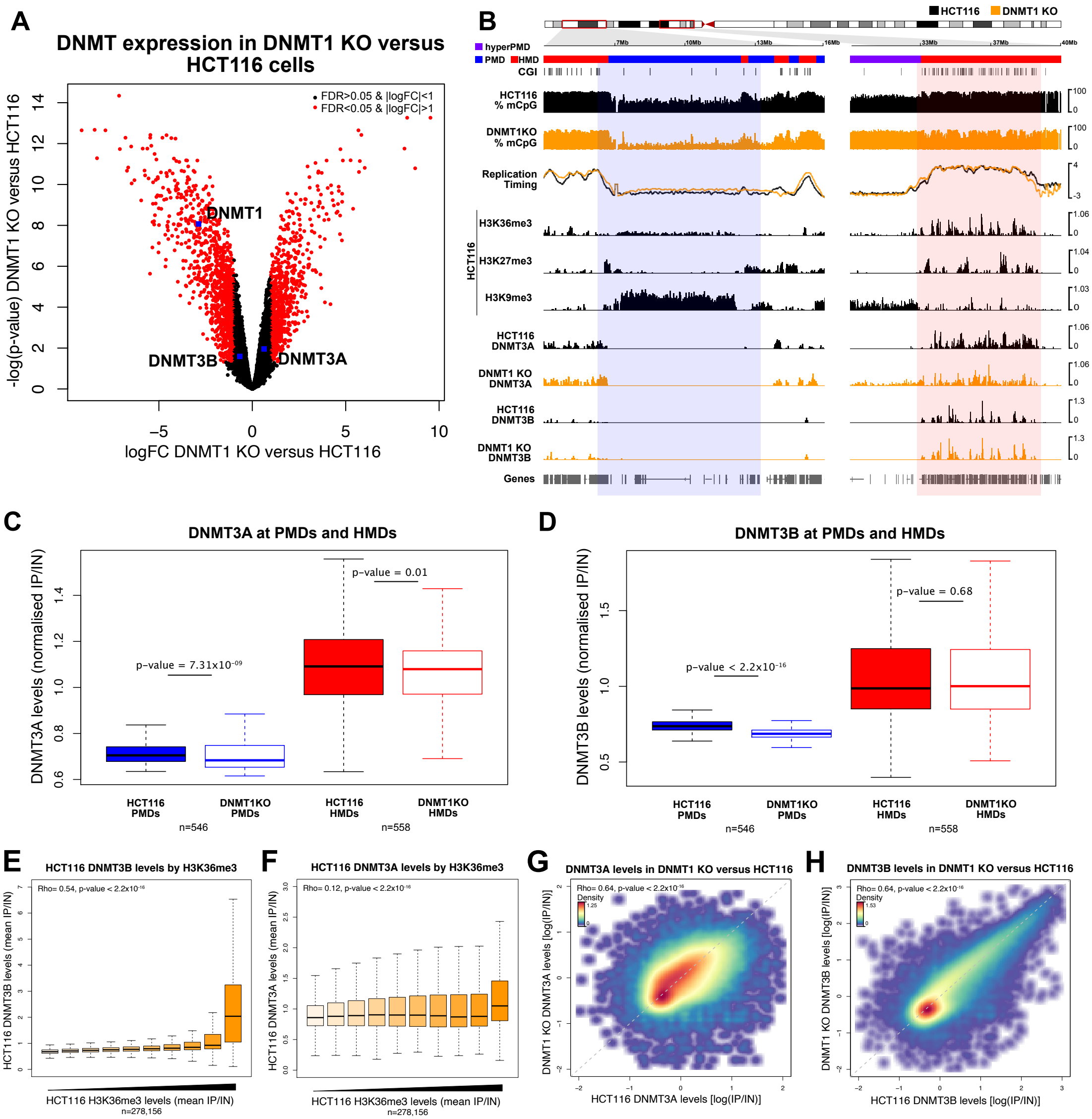

A
DNMT expression in DNMT1 KO versus HCT116 cells
FDR>0.05 & |logFC|<1
FDR<0.05 & |logFC|>1
DNMT1
DNMT3B
DNMT3A
-log(p-value) DNMT1 KO versus HCT116
logFC DNMT1 KO versus HCT116
B
HCT116
DNMT1 KO
hyperPMD
PMD
HMD
CGI
HCT116 % mCpG
DNMT1KO % mCpG
Replication Timing
HCT116
H3K36me3
H3K27me3
H3K9me3
HCT116 DNMT3A
DNMT1 KO DNMT3A
HCT116 DNMT3B
DNMT1 KO DNMT3B
Genes
C
DNMT3A at PMDs and HMDs
DNMT3A levels (normalised IP/IN)
p-value = 7.31x10-09
p-value = 0.01
HCT116 PMDs
DNMT1KO PMDs
HCT116 HMDs
DNMT1KO HMDs
n=546
n=558
D
DNMT3B at PMDs and HMDs
DNMT3B levels (normalised IP/IN)
p-value < 2.2x10-16
p-value = 0.68
HCT116 PMDs
DNMT1KO PMDs
HCT116 HMDs
DNMT1KO HMDs
n=546
n=558
E
HCT116 DNMT3B levels by H3K36me3
Rho= 0.54, p-value < 2.2x10-16
HCT116 DNMT3B levels (mean IP/IN)
HCT116 H3K36me3 levels (mean IP/IN)
n=278,156
F
HCT116 DNMT3A levels by H3K36me3
Rho= 0.12, p-value < 2.2x10-16
HCT116 DNMT3A levels (mean IP/IN)
HCT116 H3K36me3 levels (mean IP/IN)
n=278,156
G
DNMT3A levels in DNMT1 KO versus HCT116
Rho= 0.64, p-value < 2.2x10-16
Density
DNMT1 KO DNMT3A levels [log(IP/IN)]
HCT116 DNMT3A levels [log(IP/IN)]
H
DNMT3B levels in DNMT1 KO versus HCT116
Rho= 0.64, p-value < 2.2x10-16
Density
DNMT1 KO DNMT3B levels [log(IP/IN)]
HCT116 DNMT3B levels [log(IP/IN)]

**Figure S4. DNMT3A localises to hypermethylated PMDs.**

(**A**) Volcano-plot showing differential expression of protein coding genes between HCT116 and DNMT1KO cells. DNMT1, DNMT3A and DNMT3B are indicated. FC = fold change. (**B**) Representative genomic loci showing DNMT3A/B localisation at a H3K9me3-marked PMD and an HMD in DNMT1 KO cells. Genome browser plots showing DNA methylation levels (mCpG) alongside DNMT3A/B ChIP-seq and HCT116 histone modifications and repli-seq. DNA methylation levels are plotted in 10 kb genomic windows. ChIP-seq tracks are normalised $\log_{10}$ IP/IN. Replication timing data are loess smoothed repli-seq early/late ratios over 10 kb. Representative H3K9me3-marked PMD and HMD are indicated by the coloured boxes. CGI = CpG islands. (**C**) Boxplot comparing DNMT3A levels at PMDs (n = 546 domains) and HMDs (n = 558 domains). ChIP-seq data are mean normalised IP/IN. P-value from two-sided Wilcoxon rank sum test. (**D**) Boxplot comparing DNMT3B levels at PMDs (n = 546 domains) and HMDs (n = 558 domains). ChIP-seq data are mean normalised IP/IN. P-value from two-sided Wilcoxon rank sum test. (**E**) Boxplot showing mean HCT116 DNMTB levels in 10 kb genomic windows divided in deciles according to their mean levels of H3K36me3 in HCT116 cells. ChIP-seq data are normalised IP/IN. Spearman's correlation coefficient (Rho) is shown alongside its associated p-value and n is the number of windows analysed. (**F**) Boxplot showing mean HCT116 DNMT3A levels in 10 kb genomic windows divided in deciles according to their mean levels of H3K36me3 in HCT116 cells. ChIP-seq data are normalised IP/IN. Spearman's correlation coefficient (Rho) is shown alongside its associated p-value and n is the number of windows analysed. (**G**) Density scatter plot showing genome-wide correlation DNMT3A levels in DNMT1 KO and HCT116 cells. ChIP-seq data are normalised log IP/IN in 10kb windows. Spearman's correlation (Rho) and associated p-value is shown. (**H**) Density scatter plot showing genome-wide correlation DNMT3B levels in DNMT1 KO and HCT116 cells. ChIP-seq data are normalised log IP/IN in 10kb windows. Spearman's correlation (Rho) and associated p-value is shown. For boxplots: Lines = median; box = 25th–75th percentile; whiskers = 1.5 × interquartile range from box. All histone ChIP-seq and repli-seq data shown are derived from the mean of two biological replicates.

### Fig. S5

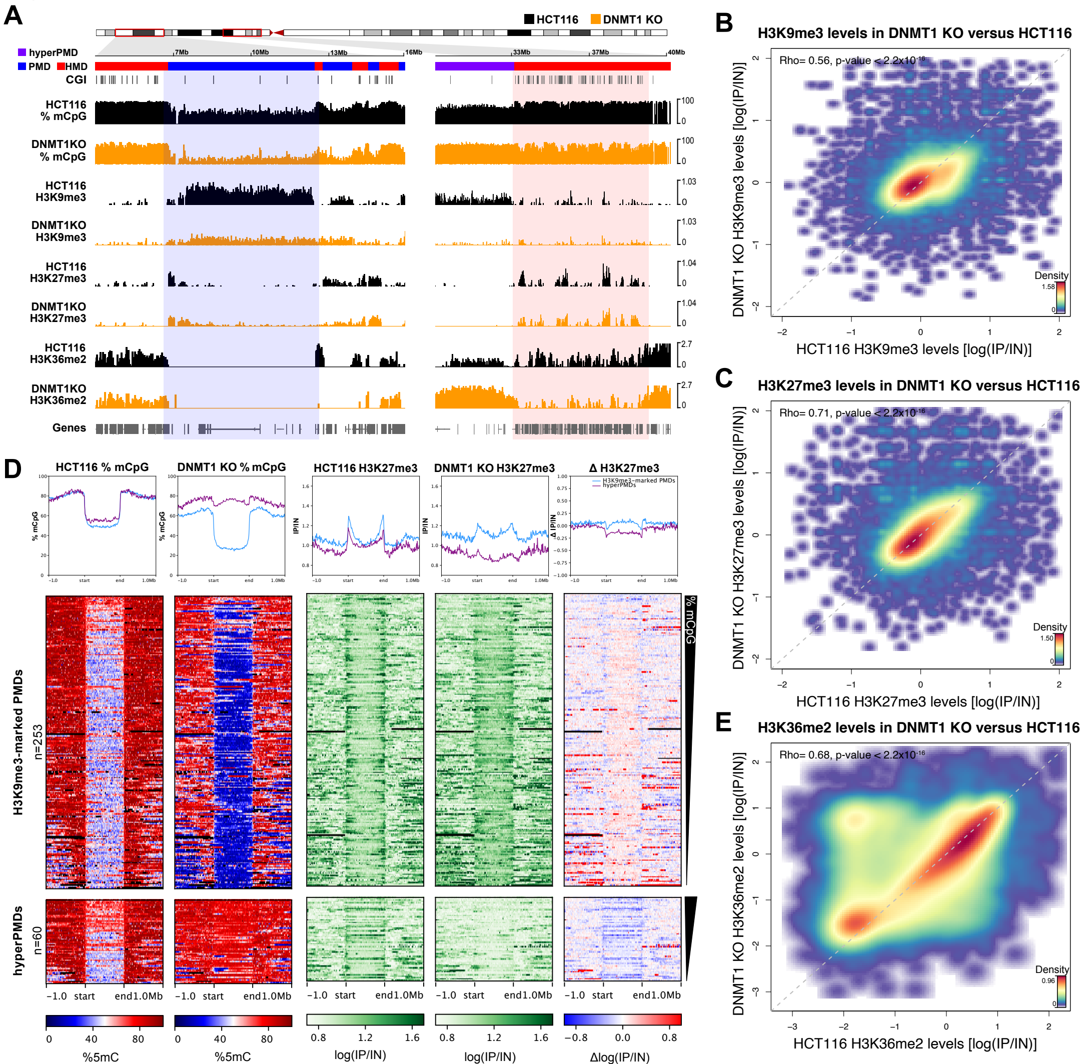
A
HCT116
DNMT1 KO
hyperPMD
PMD
HMD
CGI
HCT116 % mCpG
DNMT1KO % mCpG
HCT116 H3K9me3
DNMT1KO H3K9me3
HCT116 H3K27me3
DNMT1KO H3K27me3
HCT116 H3K36me2
DNMT1KO H3K36me2
Genes
7Mb
10Mb
13Mb
16Mb
33Mb
37Mb
40Mb
B
H3K9me3 levels in DNMT1 KO versus HCT116
Rho= 0.56, p-value < 2.2x10-16
DNMT1 KO H3K9me3 levels [log(IP/IN)]
HCT116 H3K9me3 levels [log(IP/IN)]
Density
C
H3K27me3 levels in DNMT1 KO versus HCT116
Rho= 0.71, p-value < 2.2x10-16
DNMT1 KO H3K27me3 levels [log(IP/IN)]
HCT116 H3K27me3 levels [log(IP/IN)]
Density
E
H3K36me2 levels in DNMT1 KO versus HCT116
Rho= 0.68, p-value < 2.2x10-16
DNMT1 KO H3K36me2 levels [log(IP/IN)]
HCT116 H3K36me2 levels [log(IP/IN)]
Density
D
HCT116 % mCpG
DNMT1 KO % mCpG
HCT116 H3K27me3
DNMT1 KO H3K27me3
Δ H3K27me3
H3K9me3-marked PMDs
hyperPMDs
% mCpG
IP/IN
Δ IP/IN
-1.0 start end 1.0Mb
H3K9me3-marked PMDs n=253
hyperPMDs n=60
%5mC
log(IP/IN)
Δlog(IP/IN)

**Figure S5. Hypermethylated PMDs lose H3K9me3 and gain H3K36me2.**

(**A**) Representative genomic loci showing H3K9me3 and H3K36me2 levels at a representative H3K9me3-marked PMD and an HMD in DNMT1 KO cells. Genome browser plots showing DNA methylation levels (mCpG) alongside H3K9me3, H3K27me3 and H3K36me2 ChIP-seq. DNA methylation levels are plotted in 10 kb genomic windows. ChIP-seq tracks are normalised $\log_{10}$ IP/IN. Representative hypermethylated PMD is indicated by the coloured box. CGI = CpG islands. (**B**) Density scatter plot showing genome-wide correlation H3K9me3 levels in DNMT1 KO and HCT116 cells. ChIP-seq data are normalised log IP/IN in 10kb windows. Spearman's correlation (Rho) and associated p-value is shown. (**C**) Density scatter plot showing genome-wide correlation H3K27me3 levels in DNMT1 KO and HCT116 cells. ChIP-seq data are normalised log IP/IN in 10kb windows. Spearman's correlation (Rho) and associated p-value is shown. (**D**) Heatmaps and pileup plots of HCT116 and DNMT1 KO DNA methylation levels alongside H3K27me3 levels for hypermethylated PMDs (n= 60) and all other H3K9me3-marked PMDs (n= 253). ChIP-seq data are mean normalised log IP/IN. DNA methylation levels are mean % mCpG. PMDs are aligned and scaled to the start and end points of each domain and ranked based on their mean methylation levels in HCT116 cells. (**E**) Density scatter plot showing genome-wide correlation H3K36me2 levels in DNMT1 KO and HCT116 cells. ChIP-seq data are normalised IP/IN in 10kb windows. Spearman's correlation (Rho) and associated p-value is shown. All histone ChIP-seq data shown are derived from the mean of two biological replicates.
